# Curvature sensing and generation by diverse membrane-tethered polymers can be explained by a unifying physical model

**DOI:** 10.64898/2026.09.28.754990

**Authors:** Ke Xiao, Feng Yuan, Wade F. Zeno, Justin R. Houser, Advika Kamatar, Jeanne C. Stachowiak, Padmini Rangamani

**Affiliations:** Department of Mechanical and Aerospace Engineering, University of California San Diego, La Jolla CA 92093, USA; Department of Molecular and Cell Biology, University of California, Berkeley, CA 94712, USA; Mork Family Department of Chemical Engineering and Materials Science, University of Southern California, Los Angeles, CA 90089, USA; Department of Biomedical Engineering, University of Texas at Austin, Austin, TX 78712, USA; Department of Pharmacology, School of Medicine, University of California San Diego, La Jolla CA 92093, USA

**Keywords:** Membrane-tethered polymers, ionic strength, degree of ionization, polymer brush theory, Helfrich energy, membrane bending, endocytosis, intrinsically disordered proteins

## Abstract

Cell membranes are decorated by dense, polymeric macromolecules such as proteins with bulky intrinsically disordered regions, glycosylated proteins, and glycopolymers. Experiments have illustrated that the curvature of cellular membranes, which plays a role in diverse biological processes, can be regulated by the local density of these membrane-tethered polymers and their response to environmental changes. However, the mechanisms underlying the feedback between solution conditions, membrane curvature, and the local density and molecular weight of membrane-tethered polymers remain largely unexplored. In this work, using a combination of theory and experiment, we show that the ionic strength of the surrounding solution and the degree of ionization both play important roles in regulating the conformation of the membrane-tethered polymers, which in turn influences their membrane curvature generation capabilities. Specifically, we show that for negatively charged, disordered proteins and glycans, the thickness of the polymer brush decreases with ionic strength but increases with grafting density and degree of ionization, resulting in enhanced sensitivity to membrane curvature and an amplified ability to drive membrane curvature at lower ionic concentrations. Our findings reveal a mechanism by which membrane-tethered polymers generate membrane curvature to minimize free energy, with the extent of curvature generation modulated by the polymer grafting density and layer thickness, as well as by the ionic strength of the surrounding solution. The implications of these findings are broad, suggesting that perturbations to biopolymer charge, such as sialylation of glycans and phosphorylation of disordered proteins, could actively modulate membrane curvature induction in diverse cellular processes from endocytosis to cell motility.

**Significance Statement:** Cellular membranes are richly decorated by dense polymeric structures, including the glycocalyx and membrane-bound proteins with bulky intrinsically disordered domains. These biopolymers often have a high net-negative charge, such that they are sensitive to changes in the ionic landscape in the surrounding solution. They respond to such changes in their environment in multiple ways including by sensing local membrane curvature, changing their organization to shield the cell, and generating curvature to facilitate intra- and intercellular communication. However, a predictive framework that can relate solution conditions to the membrane-tethered polymer properties and in turn to their curvature generating abilities remains lacking. To address this gap, we present a unifying biophysical model that provides a physical explanation of how changes in the solution environment modulate interactions between polymer chains within the brush layer. Our findings predict that tuning the ionic strength of the extracellular space, along with the properties of the membrane-tethered polymers, can significantly modulate curvature sensing and curvature generation. These findings provide a predictive framework for understanding the role of self-avoiding biopolymers during diverse cellular processes from endocytosis to cell motility.

## Introduction

The surface of cellular membranes contains a large number of polymeric macromolecules including peripheral proteins, glycosylated proteins, glycolipids, proteoglycans, polysaccharides, and glycosaminoglycans that extend above the plasma membrane (*1*). The majority of polymeric glycoproteins and glycans are densely coated on the membrane and form a sugar-rich layer of brush-like polymer meshwork termed the glycocalyx (*2*–*5*) (Fig. 1A). These polymer-like macromolecules play an essential role in a range of cellular processes in health and disease (*5*–*9*). Another class of large membrane-tethered molecules are intrinsically disordered proteins, which are also highly abundant on cell membranes (*10*–*13*). The function of these molecules on the membrane has been investigated in different experimental studies and we now know that many of them can sense and generate membrane curvature (*14*–*17*) (Figs. 1B and 1C). Recent experiments also suggest that the hydrodynamic radius of these proteins can be modulated in response to changes in solution conditions (*14*). These bulky polymers occupy a large volume on the membrane surface. Steric pressure between them can drive membrane curvature, and, correspondingly, the constraints placed on their conformational entropy by the membrane surface give rise to their documented ability to sense membrane curvature (*4, 16*).

**Figure 1:**
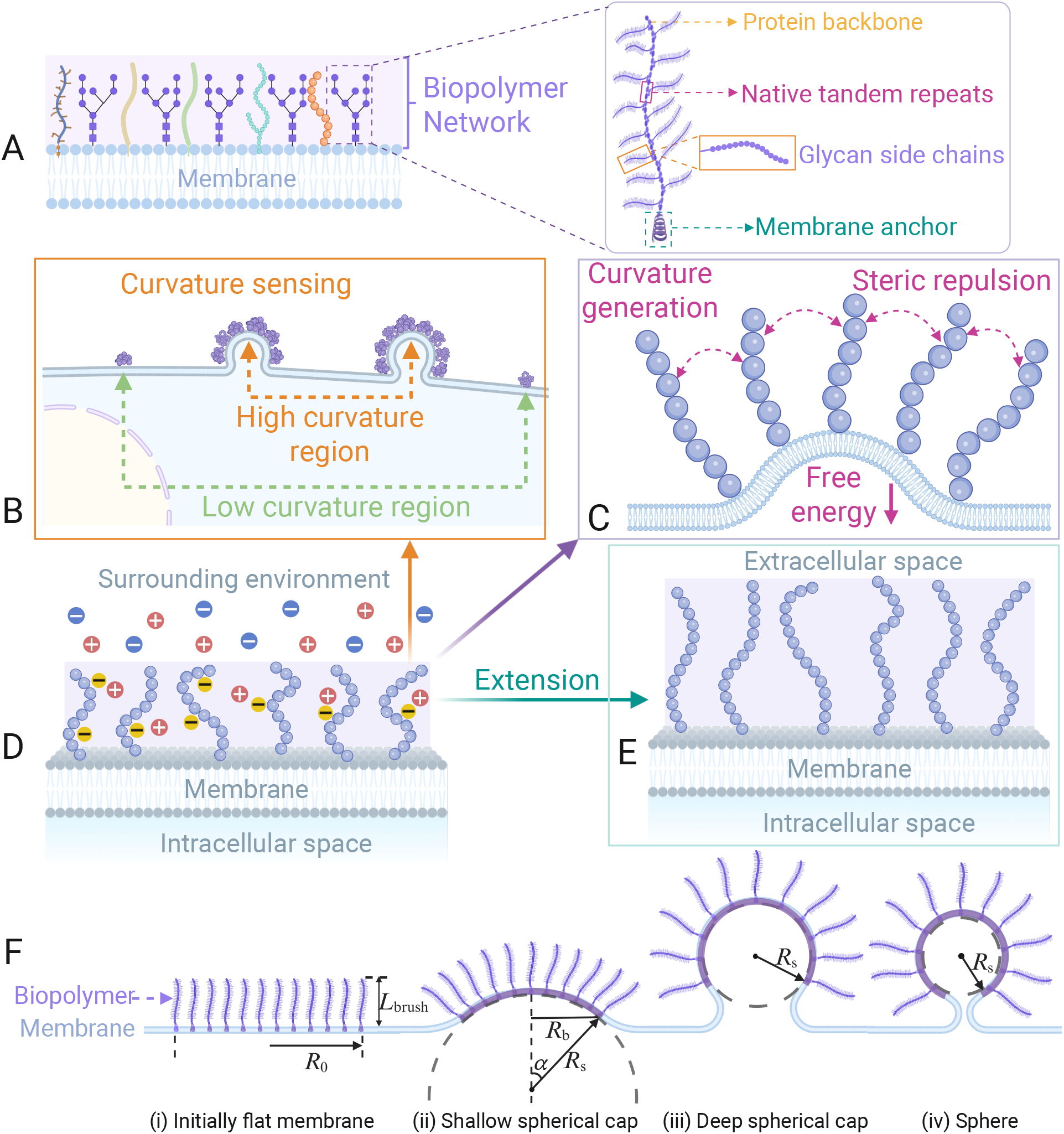
Interaction between membrane-tethered polymers and the lipid bilayer. (A) Schematic showing the crowded biopolymer network on a cell membrane such as a sugar-rich layer of glycocalyx. An enlarged view of the polymer structure of glycocalyx constituents such as mucins. Schematic representation of membrane-tethered polymers (B) sensing local membrane curvature and (C) generating membrane curvature. (D) Schematic representation of a cell membrane decorated with a polymer brush network immersed in an aqueous environment. Typically, membrane-tethered polymers are negatively charged and are surrounded by mobile ions in the salt solution. (E) Schematic illustration of the height regulation of membrane-tethered polymers. (F) Mechanism of biopolymer-mediated membrane bending. The polymers are modeled as grafted on a flat membrane patch of given area with radius *R*_0_, and repulsive interactions between the polymers can lead to membrane curvature. In our model, the membrane consists of two parts: a patch domain with membrane-tethered polymers (purple) and the remaining free membrane region.

Another important feature of these membrane-tethered polymers is that they are typically charged (Fig. 1D), and have electrostatic properties. For example, the dissociated acidic groups of sialic acid within the glycocalyx make it a negatively charged polymer brush, which leads to electrostatic interactions (*2*). Mucins are estimated to carry approximately five electron charges per 20 amino acid tandem repeat regions (*4*). Many disordered proteins such as AP180 also have a substantial net negative charge owing to negatively charged amino acids (*18*). Thus, the electrostatic properties of these macromolecules are directly associated with the degree of ionization of their acidic groups. The pH of the surrounding extracellular solution will also have an impact on the degree of ionization of these membrane-bound polymers. As a consequence, changes in salt concentration or pH can tune the molecular interactions between these membrane-bound moieties. Interestingly, the ionic strength also affects the length of biopolymers themselves (*19*–*21*). Together, the degree of ionization (or pH) and ionic strength are crucial factors for the cell in regulating biomolecular interactions.

For a cell membrane coated with a polymer brush network, the steric effect between polymer chains in the crowded environment can alter the height of these polymers (Fig. 1E). Here, the height is defined as the distance between the membrane surface and the portion of the polymer that is furthest from the surface. If a membrane-tethered polymer experiences lateral pressure, which may arise from other biomolecules crowding the same membrane surface, it tends to adopt a more extended conformation, such that the height increases. This height regulation is important for cell signaling (*7, 8*), pathogen defense (*22*), and cancer development (*6, 23*). Using cell surface optical profilometry, recent studies have revealed that the average height of multidomain proteins on cell surfaces strongly depends on surface and solution crowding (*24, 25*). In addition, previous work has shown that DNA conformation is strongly regulated by its surrounding environment (*20, 21*), including ionic strength and the crowding of biopolymers. Collectively, all these experimental studies indicate that a wide variety of membrane-tethered polymers can sense and generate membrane curvature as a function of grafting density and height, modulate their brush height in response to the solution environment, which further reinforces their curvature generation capabilities. Here, we seek to ask the question: are there any unifying physical principles that can explain these disparate observations? To answer this question, we developed a theoretical biophysical model.

An extensive body of theoretical studies has been devoted to understanding the shape changes of membranes arising from the polymer chains anchored to membrane surfaces (*26*–*34*). Previous studies using analytical methods, scaling theory, and Monte Carlo simulations have shown that polymers grafted onto one side of the plasma membrane can lead to changes in membrane shape and local inhomogeneities of bending rigidity and spontaneous curvature of the membrane (*26*– *32*). Moreover, the concentration gradient of anchored polymers and the concentration of homogeneous anchored amphiphilic polymers have been shown to govern the formation of membrane tubes and pearling instabilities (*33, 34*). Combining the Helfrich curvature elasticity theory with polymer physics for the glycocalyx has previously allowed us to explore the collective effects of biopolymers on membrane shape transformation (*16, 35, 36*). Although previous models have examined membrane remodeling driven by polymer crowding and membrane bending, they have focused on specific experimental observations rather than explaining multiple disparate observations. As a result, a unified theoretical framework that simultaneously accounts for polymer crowding, membrane bending elasticity, and ionic effects remains lacking. In this work, we close this gap and present a unifying framework to investigate the relationship between the ionic strength, membrane curvature, and height of membrane-tethered polymers on cell membranes.

## Model Development and Methods

In this section, we provide a brief overview of the theory for membrane curvature sensing and membrane curvature generation. We assume that the system is in mechanical equilibrium at all times, implying that the system selects the membrane shape that minimizes the total free energy of the composite system. Detailed derivations are provided in the Supporting Information (SI).

### I. Membrane Curvature Sensing by Biopolymers

Membrane-tethered polymers have recently been discovered to exhibit curvature-sensing behavior (*14*–*16*). To understand the interaction between ionic strength, distribution of membrane-tethered polymers, and curvature sensing, we developed a thermodynamic model. The free energy of the system can be written as (*37*–*41*)

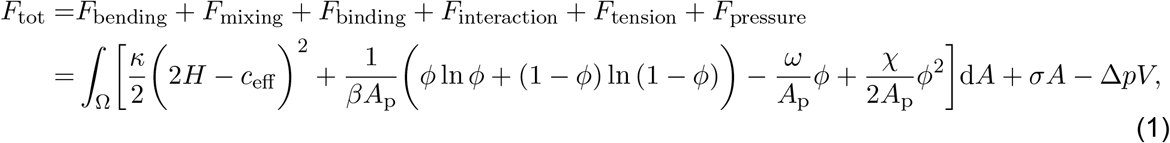

where *κ* is the bending modulus of the lipid bilayer, *H* = (*c*_1_ + *c*_2_)*/*2 is the mean curvature in which *c*_1_ and *c*_2_ are the two principal curvatures of the local membrane surface, *c*_eff_ is the effective spontaneous curvature of the membrane. We assume that this spontaneous curvature is generated due to the steric interactions of the membrane-tethered polymers. *β* = 1*/*(*k*_*B*_*T*) with *k*_*B*_ being the Boltzmann’s constant and *T* being the temperature, *A*_p_ is the surface area occupied by the anchor of the polymer, *ϕ* is the area fraction of the polymers on the membrane, *ω* represents the chemical potential of tethering, *χ* is a direct interaction coefficient between the polymers (*χ >* 0 and *χ <* 0 represent repulsive and attractive interactions respectively). *σ* is the membrane tension, which acts as a Lagrange multiplier to keep the total membrane area *A* constant, and Δ*p* is the hydrostatic pressure difference across the cell membrane.

To quantify the spatial heterogeneity of these membrane-tethered polymers on cell surfaces with different local membrane curvature, we consider the polymer-membrane system consisting of a vesicle component that represents the whole cell and a long membrane tether connected to the cell body, as shown in Fig. 2A. The difference between the area fraction of polymers on the membrane tube *ϕ*_t_ and on the planar cell surface *ϕ*_V_ is given by Δ*ϕ* = *ϕ*_t_ − *ϕ*_V_. By minimizing the total free energy in Eq. (1), we obtain the relationship between the relative partitioning number 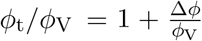 and the curvature-sensing capabilities of these membrane-tethered moieties. A detailed description is provided in the SI Appendix, section 1.

**Figure 2:**
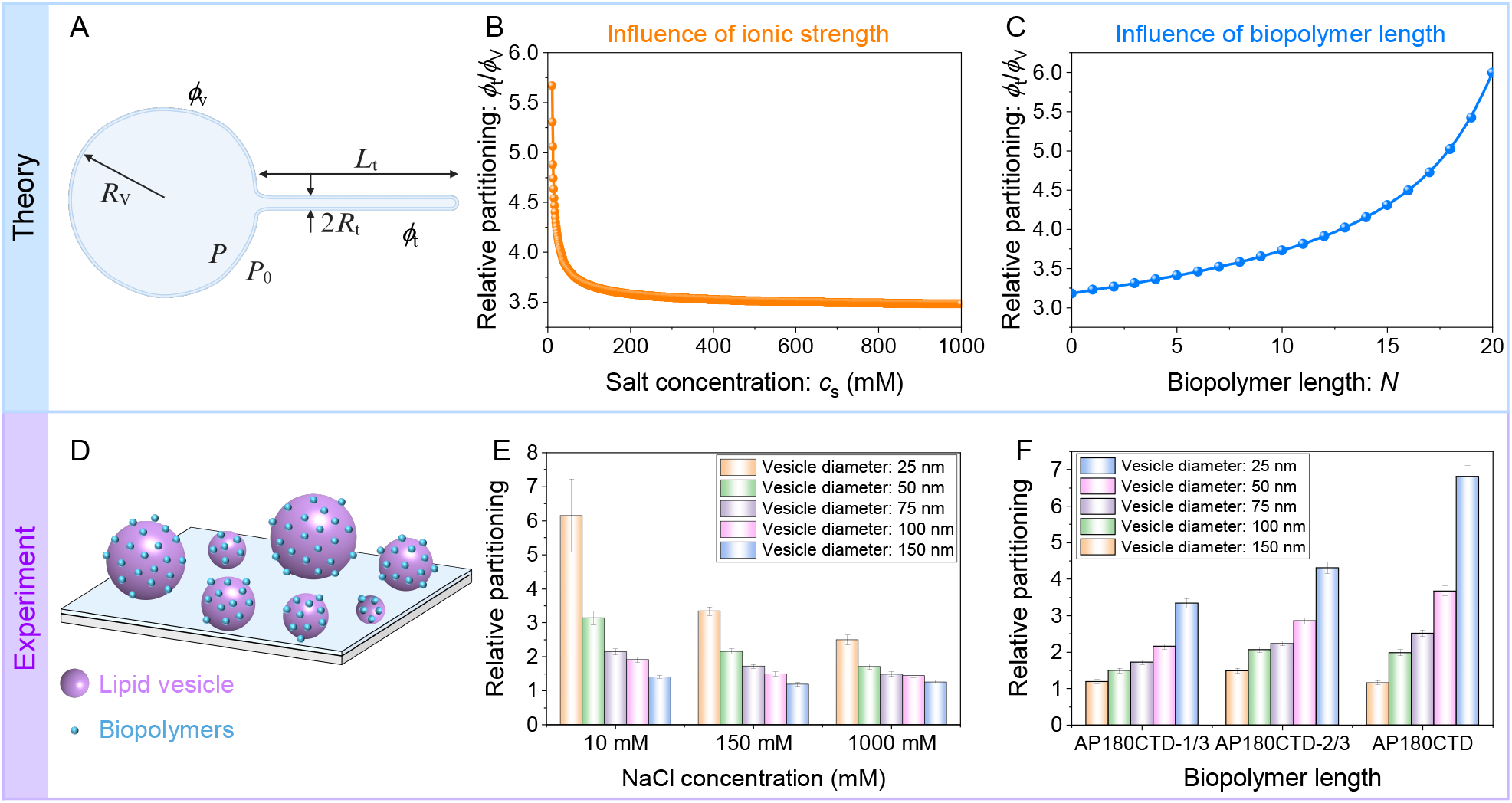
Curvature sensing of membrane-tethered polymers. (A) A schematic illustration of the system geometry used in the curvature-sensing model. Theoretical prediction of the dependence of relative partitioning on (B) ionic strength and (C) biopolymer length. (D) Schematic of the tethered vesicle assay utilized to quantify the curvature-sensing properties. (E) Relative partitioning of the AP180CTD-1/3 protein as a function of sodium chloride concentration. (F) Bar plots showing relative partitioning for different vesicle sizes and various AP180CTD truncation mutants (AP180CTD-1/3, AP180CTD-2/3, and AP180CTD), all of which are polymer-like disordered domains. The experimental data shown in (E) and (F) are originally from Ref. (*14*) and reanalyzed for the current work.

### II. Membrane Curvature Generation and Modulation of Brush Height

Next, we developed a model for membrane-curvature generation that takes into account the ionic environment (Fig. 1F). We assumed that the initial state of the membrane is flat; this patch undergoes deformation in response to the stresses exerted by the polymer layer. We modeled the membrane-tethered polymers as a layer of uniform height, known as a polymer brush-like structure, where the polymer brushes are extended equally in height, *L*_brush_ (see Fig. 1F). For simplicity, we model the grafted region as portions of spherical caps. As a result, the membrane is divided into two parts: (i) the membrane patch grafted with polymers (spherical cap part with radius *R*_s_ and opening angle *α*); and (ii) the free part of the membrane. We also assume rotational symmetry of the membrane surface with respect to the *z* axis.

Following the classical Canham–Helfrich continuum membrane model and combining the polymer brush theory, the total free energy of the polymer-membrane composite system comprises four terms

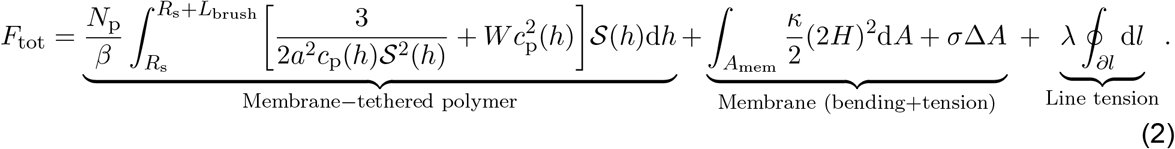

The first term is the energy contribution associated with the membrane-tethered polymers, which includes the elastic stretching of the polymer chains, the interactions between monomers in the polymer brush, and the electrostatic contribution for ionic brushes. Here *N*_p_ = *ρA*_0_ is the number of polymer chains grafted on the membrane with *ρ* and *A*_0_ being the grafting density and the membrane patch area covered by biopolymers, *a* is the effective monomeric segment size, *c*_p_(*h*) is the local monomer density profile along the thickness with *h* being the radial distance defined from the center of the spherical surface, *S*(*h*) is the area per chain at radial distance *h* from the center of the spherical membrane surface, 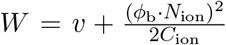 with *v, ϕ*_b_, *N*_ion_, and *C*_ion_ being the second virial coefficient, the degree of ionization, the number of ionizable sites, and the ion concentration, respectively. Note that the brush height *L*_brush_ is obtained from the conservation condition and is a function of *C*_ion_ and *ϕ*_b_ (see Eq. (S39)). Consequently, the height of membrane-tethered polymers changes with the variation of ionic strength and the degree of ionization at flat and curved membrane surfaces (see Figs. S5 and S6 for the dependence of brush height on *C*_ion_ and *ϕ*_b_ at flat and curved membrane surfaces). The second bending energy term and the third surface tension energy term that appear in Eq. (2) are the elastic energy of the cell membrane, where Δ*A* is the increase of membrane area (from flat case) due to the deformation of membrane shape. The last term on the right-hand side of Eq. (2) represents the interfacial energy arising from line tension at the boundary between the polymer-grafted and ungrafted regions. Here, *λ* is the line tension strength, and the integral is taken along the domain periphery, d*l*. The detailed derivation of each individual energy component in Eq. (2) is provided in SI Appendix, sections 2–3. Using these expressions, we can eventually split the total free energy into two parts: the contribution of the membrane patch covered with polymer and the contribution from the free part of the membrane, i.e., 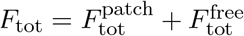. Here, the first term reads

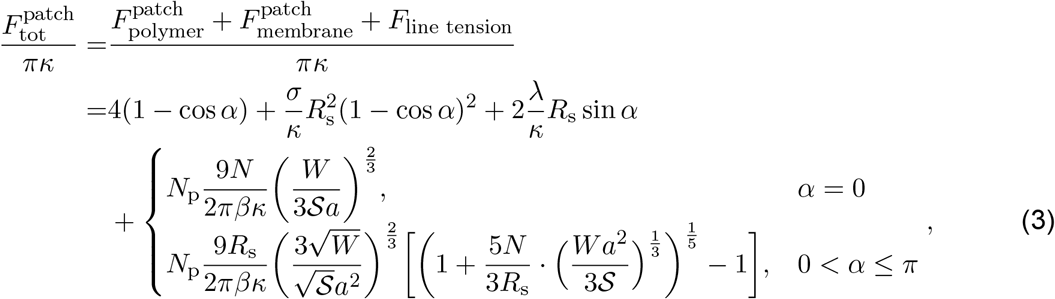

where *N* is the number of segments per polymer chain. For the free part of the membrane (outside the polymer-grafted region), we assume the membrane is axisymmetric and is parametrized by *r*(*s*), *z*(*s*) and *ψ*(*s*) (see Fig. S7 in SI Appendix, Section 4), where *s* is the arc length of the free membrane. Therefore, the total energy of the free membrane can be written as 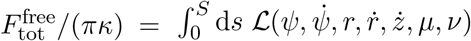 (*42, 43*), where *ℒ* is a Lagrangian defined by

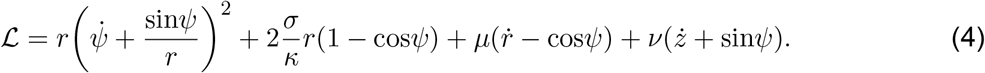

Here, the dots denote a derivative with respect to the arc length, and *µ*(*s*) and *ν*(*s*) are Lagrange multipliers to impose the geometric constraints between *r, z* and *ψ*. A variation of the energy functional of the free membrane against the shape variables *r*(*s*), *z*(*s*), and *ψ*(*s*) produces a set of shape equations, the derivation of which can be found in SI Appendix, section 4.

### III. Numerical Implementation

For the curvature-sensing model, we evaluated the area fraction of membrane-tethered polymers on the cell surface to quantify the curvature sensing ability under varying ionic strengths and polymer lengths. To determine the brush height, we first compute the equilibrium membrane shape by minimizing the total free energy and then calculate the height using Eq. (S39). In general, the shape equations have to be solved numerically due to their nonlinearity. Together with the corresponding boundary conditions, the membrane shape profile and its corresponding total energy *F*_tot_(*α*) are obtained numerically for different values of the opening angle *α* under the constraint of fixed membrane patch area 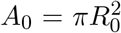. Based on the optimal opening angle *α*_min_ obtained via minimizing the total energy, we can determine the equilibrium geometry of the membrane. The values of parameters used in the model are summarized in Table S1 unless otherwise specified.

## Results

### Membrane curvature sensing by biopolymers

The preferential localization of biopolymers on regions of higher local membrane curvature is referred to as curvature sensing. To understand how charged, membrane-bound biopolymers sense membrane curvature, we developed a thermodynamic model for the heterogeneity of biopolymer distribution as a function of membrane curvature (see SI Appendix, section 1). Specifically, we define a membrane surface consisting of a cylindrical membrane tube surrounded by a nearly flat lipid membrane. In this model, we defined a relative partitioning number, *ϕ*_t_*/ϕ*_V_, to quantify the curvature-sensing ability of biopolymers. Here, *ϕ*_t_ is the area fraction of biopolymers on the membrane tube, and *ϕ*_V_ is the area fraction on the planar cell surface (see Fig. 2A). We calculate the relative partitioning number as a function of ionic strength and biopolymer length in Figs. 2B and 2C, respectively. Our model predicts a decrease in relative partitioning with increasing ionic strength (Fig. 2B) but an increase with increasing biopolymer length (Fig. 2C). These trends indicate that biopolymers preferentially partition to higher membrane curvature regions and that this curvature-sensing ability is modulated by both the ionic strength of the solution and biopolymer length. The mechanism underlying this ionic strength-dependent curvature-sensing behavior is attributed to electrostatic effects (*14, 44*). At low ionic strength, strong electrostatic effect enlarges the effective biopolymer size and enhances repulsive interactions between polymer chains. As a result, to alleviate excluded-volume interactions and maximize available space, the chains preferentially partition toward regions of high membrane curvature. At high ionic strength, by contrast, screening of electrostatic effects diminishes this curvature sensitivity.

To test this prediction, we used the endocytic protein AP180 to examine how its curvature sensing behavior depends on salt concentration. AP180 contains a long intrinsically disordered C-terminal domain (AP180CTD) (*17*), which provides a useful model for investigating how polymer-like disordered proteins interact with membrane surfaces and sense membrane curvature. To further explore how the curvature sensitivity of this disordered domain depends on chain length, we created three AP180CTD constructs with different lengths corresponding to the full-length disordered domain, the first two-thirds (AP180CTD-2/3), and the first one-third (AP180CTD-1/3). The curvature-sensing properties of each truncation of AP180CTD were quantified by employing the tethered vesicle assay depicted in Fig. 2D (refer to Ref. (*14*) for detailed descriptions). By measuring relative protein binding to substrate-tethered vesicles of increasing diameter, we evaluated curvature-sensing behavior under various conditions (salt concentrations and chain lengths), as shown in Figs. 2E and 2F. By quantitatively comparing curvature sensing by AP180CTD-1/3 at different NaCl concentrations, Fig. 2E reveals two main results. First, the overall trend shows a substantial reduction in relative partitioning as the ionic strength of the solution increases. Second, at a fixed NaCl concentration, increasing local membrane curvature raises the relative partitioning, indicating that biopolymers preferentially bind to regions of high membrane curvature. These results demonstrate that biopolymers exhibit significant curvature sensitivity and this sensitivity is reduced with increasing ionic strength. In contrast, a direct comparison of the relative partitioning for different AP180CTD truncation mutants shows an increase in curvature sensitivity with increasing polymer chain length (Fig. 2F). For each truncation mutant, we also observed that the relative partitioning increases monotonically as membrane curvature increases (Fig. 2F). Note that AP180CTD and its truncation mutants are polymer-like disordered domains. Therefore, these data suggest that biopolymers are sensitive to membrane curvature, and this sensitivity is substantially enhanced by decreasing ionic strength and increasing biopolymer length.

### Grafting density- and length-dependent curvature generation

Thus far, we have only considered the ability of membrane-tethered polymers to sense curvature. Curvature sensing is closely related to curvature generation as both arise from the interplay between polymer steric interactions and membrane geometry. While curvature sensing describes how polymers respond to pre-existing membrane curvature, polymer crowding can in turn generate curvature. This coupling suggests a potential feedback between polymer localization and membrane deformation. However, for computational simplicity, we consider these two processes separately here. We first focus on the effects of the intrinsic physical properties of membrane-tethered polymers, specifically the grafting density *ρ* and chain length *N*, without considering any ionic effects. In these calculations, the extent of membrane deformation by the polymers is characterized by the opening angle *α*, which thus serves as a quantitative measure of curvature generation capacity. For example, *α* = 0 corresponds to a completely flattened membrane, while *α* = *π* corresponds to a sphere membrane shape. When 0 *< α < π*, the membrane with tethered polymers takes the shape of spherical caps. To assess how curvature generation depends on physical properties of membrane-tethered polymers, we performed two sets of simulations across a range of *ρ* and *N*, to determine the equilibrium membrane shape through total free energy minimization in each case. First, we plot the total energy as a function of the opening angle *α* for a series of fixed grafting densities *ρ* and polymer lengths *N*, as shown in Figs. 3A and 3B. The energy landscapes demonstrate that there exists a global minimum (blue solid dot) and a local minimum (magenta pentagon), corresponding to the stable state and metastable state, respectively. Under conditions of very low grafting density or short polymer length (e.g. *ρ* = 0.001 nm^−2^ or *N* = 10), the global minimum appears at small *α*, corresponding to a shallow dome-shaped membrane. The *α* value at this stable state defines the optimal opening angle, *α*_min_. As *ρ* or *N* increases (e.g. *ρ* = 0.002 →0.004 nm^−2^ or *N* = 15 → 20), the stable state shifts gradually, and the corresponding membrane shape evolves progressively from a shallow spherical cap to a U-shaped spherical cap. Upon increasing the grafting density or polymer length to a critical value *ρ*_c_ or *N*_c_, we observe that the total energy of the global minimum state becomes equal to that of the local minimum state, indicating that the two states coexist. Therefore, for *ρ < ρ*_c_ or *N < N*_c_, our simulation results show that the total free energy is a global minimum at 0 *< α/π <* 1, where the corresponding membrane shape is a spherical cap (see the membrane shape profiles in Figs. 3A and 3B). Further increasing grafting density or polymer length (i.e. *ρ > ρ*_c_ or *N > N*_c_) leads to the transition of membrane geometry from a spherical cap to a full sphere structure. This trend implies that a higher grafting density or longer membrane-tethered polymer on the cell membrane is necessary to achieve a high membrane curvature structure. The membrane shape exhibited under various grafting densities and polymer lengths is determined by the competition among the elastic energy (consisting of bending energy and tension) and the energy contribution associated with membrane-tethered polymers. Formation of a highly curved membrane surface requires overcoming an energetic barrier (*45*–*47*). Increasing the grafting density or polymer length can facilitate crossing this barrier, thereby enhancing curvature-generating effect and resulting in the formation of high membrane curvature shapes.

**Figure 3:**
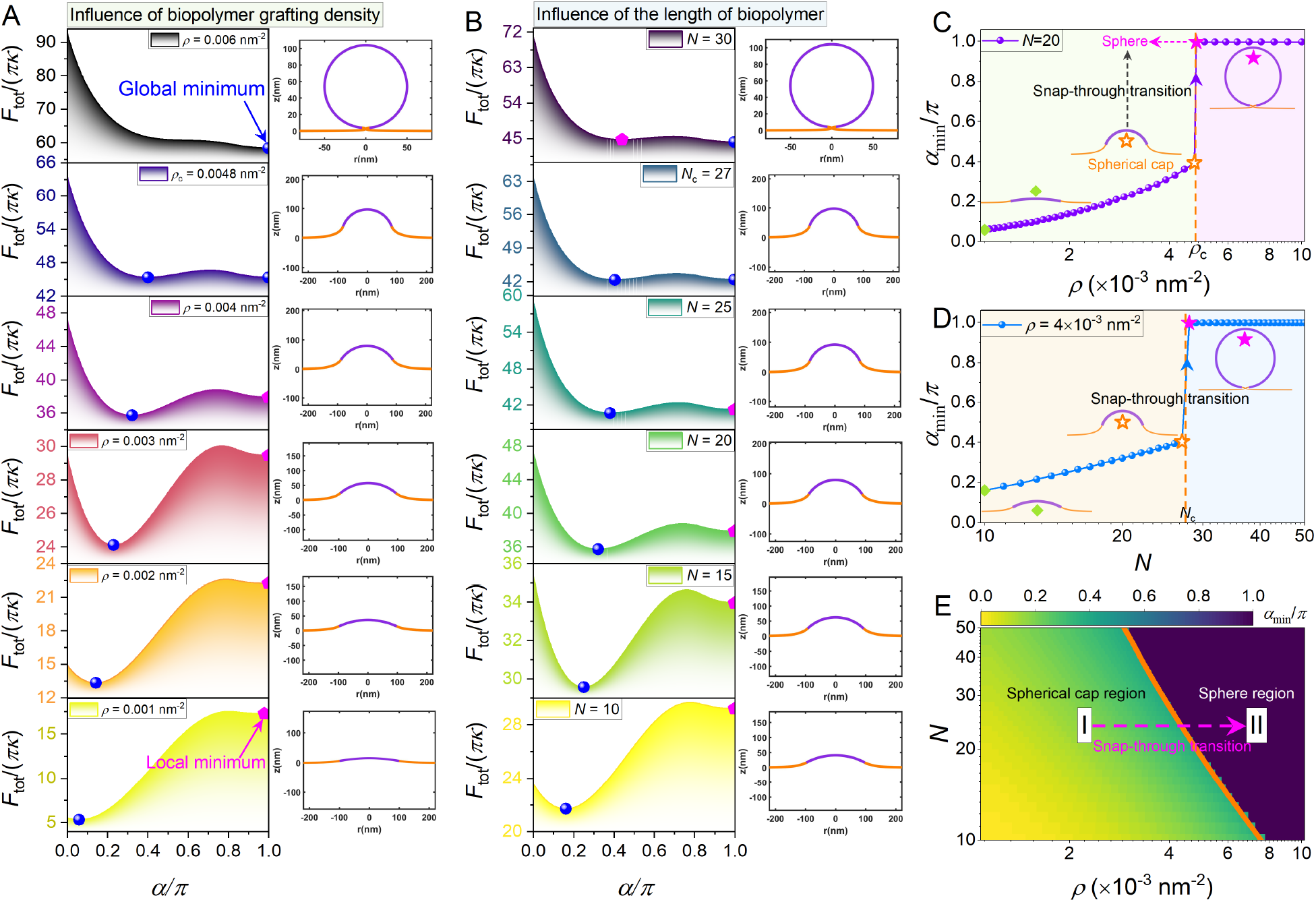
Effects of membrane-tethered polymer density and length on membrane curvature generation. Total free energy landscapes as a function of *α* for different values of membrane-tethered polymer (A) grafting density *ρ* at *N* = 20 and (B) length *N* at *ρ* = 4 *×* 10^−3^ nm^−2^. Corresponding profile views of membrane morphologies on the right panel are generated by simulations for various values of (A) *ρ* and (B) *N*. The part of the membrane grafted by polymers is in purple. The free part of the membrane is in orange. The dependence of the optimal opening angle on the (C) density and (D) length of membrane-tethered polymers. (E) Phase diagram presents the optimal opening angle (membrane morphology) as a function of grafting density *ρ* and polymer length *N*. Here, we set *ϕ*_b_ = 0 to exclude the effect of ionic strength and the other parameters are given in Table S1.

To better visualize how the properties of membrane-tethered polymers affect the membrane profile transition, we then plotted the optimal opening angle *α*_min_ as a function of grafting density and polymer length in Figs. 3C and 3D. For grafting densities or polymer lengths below the critical values *ρ*_c_ or *N*_c_, incremental increases in *ρ* and *N* induce a gradual and continuous evolution of *α*_min_ (the membrane profile). At the critical threshold, however, *α*_min_ exhibits a sharp jump, a characteristic of first-order transitions. This abrupt jump is known as a snap-through transition, which is associated with a discontinuous morphological change from a spherical cap (empty star) to a full sphere (solid star) geometry. Additionally, a logarithmic linear relationship between *α*_min_ and *N* is observed prior to this snap-through transition. To systematically map the interplay between the grafting density and polymer length in curvature generation, we further construct a two-dimensional phase diagram in the *ρ* − *N* plane (Fig. 3E), which summarizes the observed membrane morphologies. Two regimes are identified and separated by an orange boundary curve: regime I (spherical cap) and regime II (full sphere), with the color scale representing the value of the optimal opening angle. Within regime I, *α*_min_ evolves continuously with increasing *ρ* and *N*, indicating a smooth morphological progression. Beyond the boundary, the transition becomes discontinuous, characterized by an abrupt jump from a spherical cap to a full sphere. Inspecting the boundary curve in Fig. 3E, we notice that varying the polymer length shifts the critical grafting density, *ρ*_c_, at which the snap-through transition occurs. More specifically, *ρ*_c_ and *N* exhibit a logarithmic linear dependence. Our findings predict that membrane-tethered polymers generate membrane curvature toward an energetically favorable state, and that the equilibrium curvature depends on both the grafting density and the polymer length.

To test our theoretical predictions, we investigated the effect of protein concentration and intrinsically disordered region (IDR) length on membrane curvature generation. We acknowledge that the model is restricted to curvature generation as parts of spheres (see Supporting Information) for computational tractability. Therefore, we restrict our comparison between modeling and experiments to qualitative statements. Giant unilamellar vesicles (GUVs) containing DGS-NTA-Ni lipids were incubated with varying concentrations (50, 100, 250, and 1000 nM) of either the full-length intrinsically disordered C-terminal domain of AP180 (his-AP180CTD) or a truncated construct comprising its first one-third (his-AP180CTD-1/3) in buffer containing 25 mM HEPES and 150 mM NaCl (pH 7.4). Both constructs featured an N-terminal histidine tag to facilitate binding to DGS-NTA-Ni lipids on the GUV surface. Under all conditions tested, protein addition resulted in rapid recruitment to the GUV membrane within minutes, followed by the emergence and elongation of outward-protruding membrane tubules (Fig. 4A, C). Consistent with previous reports, these tubules were diffraction-limited in diameter and frequently attained lengths comparable to or exceeding the initial GUV diameter (*44*). Signal co-localization in both the protein (Atto 488) and lipid (Texas Red) fluorescence channels confirmed the presence of both components within the tubular structures (Fig. 4A, C). Quantitative analysis demonstrated that increasing protein concentration significantly elevated the frequency of outward tubule formation for both constructs (Fig. 4B, D). However, his-AP180CTD-1/3 exhibited a reduced capacity to drive membrane tubulation relative to full-length his-AP180CTD across all tested concentrations, aligning with earlier reports that the full-length domain generates stronger membrane-bending activity than truncated variants (*14, 44*). Overall, these experimental results are qualitatively consistent with our model predictions.

**Figure 4:**
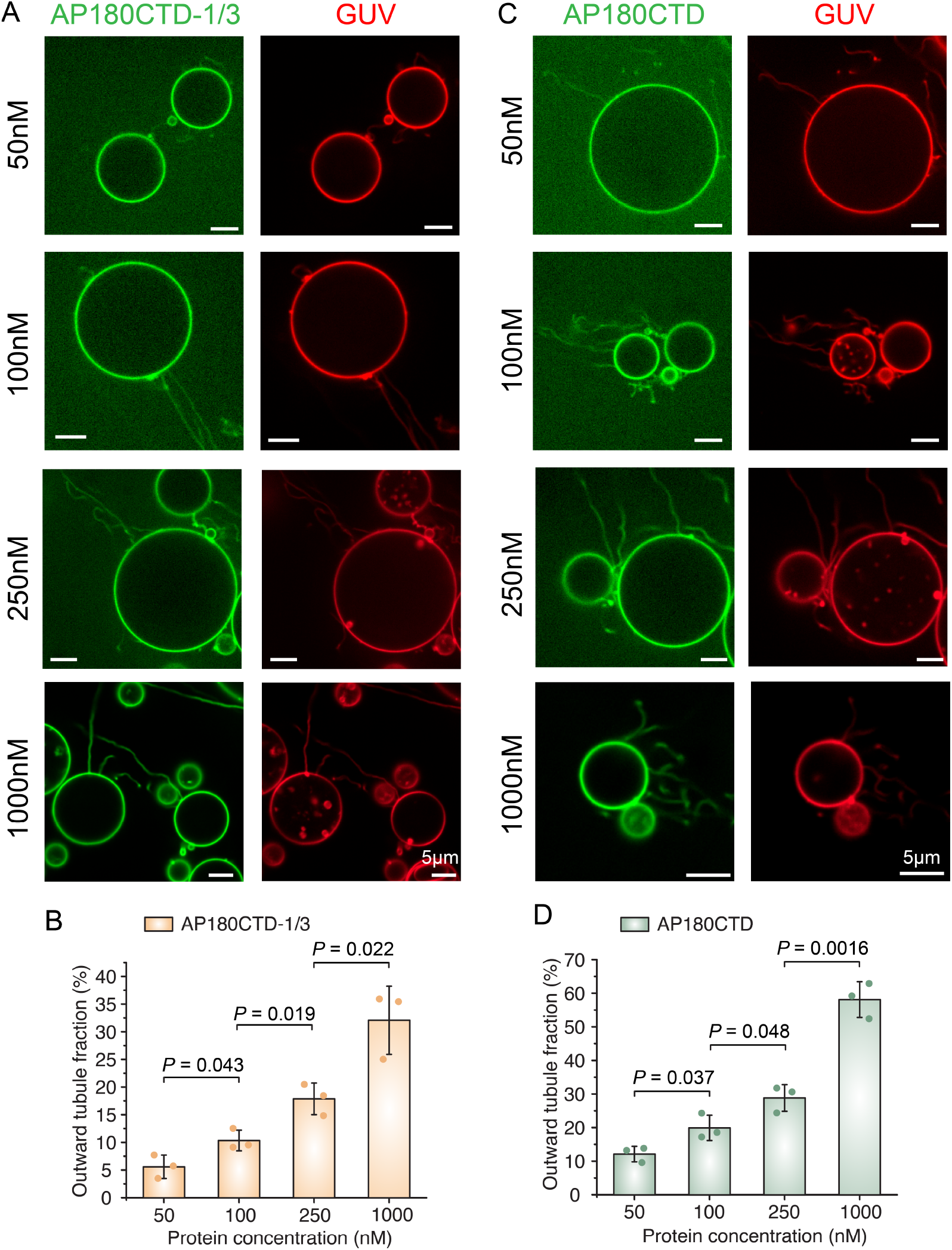
GUV tubulation as a function of protein concentration. (A, C) Representative fluorescence images of GUVs incubated with the 1/3 truncation of his-AP180CTD (A) and his-AP180CTD (C) at varying protein concentrations (50, 100, 250, and 1000 nM). Scale bar = 5 *µ*m. (B, D) Quantification of the fraction of GUVs displaying outward tubules as a function of protein concentration for his-AP180CTD-1/3 (B) and his-AP180CTD (D). All experiments were conducted in 25 mM HEPES, 150 mM NaCl buffer at pH 7.4. Error bars represent the standard deviation (SD) of three independent trials (individual data points shown). Statistical significance was evaluated using an unpaired, two-tailed Student’s t-test.

### Effects of ionization and ionic strength on membrane curvature generation

Next, we investigate the role of pH and the solution ionic strength on membrane curvature generation. We perform two series of simulations in which we keep grafting density and polymer length constant while increasing the degree of ionization *ϕ*_b_ and ionic strength *C*_ion_, respectively, allowing us to dissect their individual effects on the system. Figures 5A and 5B show plots of the total free energy landscape as a function of *α* for an increasing sequence of *ϕ*_b_ and *C*_ion_, respectively. We first observe that the energy landscapes have a global minimum (stable state, see the solid sphere dot) and a local minimum (metastable state, see the solid pentagon) that are separated by energy barriers. For degree of ionization 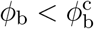, *F*_tot_(*α*) has a global minimum at 0 *< α/π <* 1, indicating that the membrane morphology regulated by the tethered polymers is a spherical cap. Increasing the degree of ionization (e.g., from *ϕ*_b_ = 0 to *ϕ*_b_ = 0.698) increases *α*_min_, which corresponds to the degree of membrane deformation (see the corresponding membrane shape on the right column in Fig. 5A). For sufficiently large *ϕ*_b_ (e.g. *ϕ*_b_ = 0.8), the global free energy minimum is shifted to a state corresponding to *α/π* = 1, which refers to a complete spherical membrane configuration.

**Figure 5:**
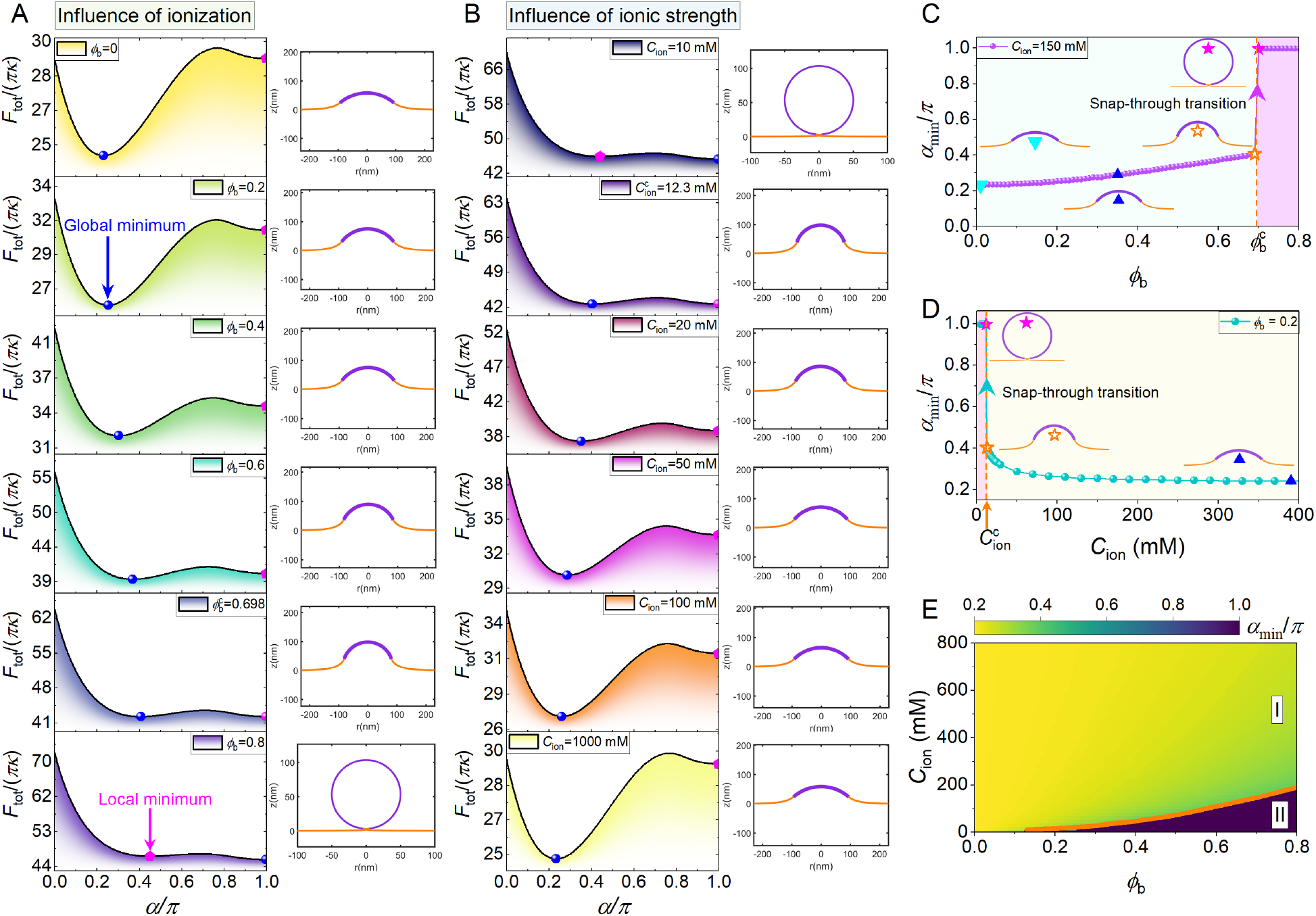
Total energy profile as a function of opening angle *α* for different (A) degree of ionization *ϕ*_b_ at *C*_ion_ = 150 mM and (B) ion concentration *C*_ion_ at *ϕ*_b_ = 0.2. The value for the other parameters are given in Table S1. Notably, the degree of ionization can be used to capture the pH of the surrounding solution because *ϕ*_b_ is related to pH (see Eq. (S33) and Fig. S3). Corresponding profile views of membrane morphologies on the right panel are generated by simulations for various values of (A) *ϕ*_b_ and (B) *C*_ion_. The dependence of normalized optimal opening angle on (C) *ϕ*_b_ and (D) *C*_ion_. (E) Two-dimensional phase diagrams on the (*ϕ*_b_ − *C*_ion_) plane characterize the interrelated effects of the degree of ionization and ionic concentration on membrane shape.

Figure 5B shows the impact of changes in ionic strength on the free energy landscape and modulation of curved membrane structures. Similar to Fig. 5A, a global energy minimum also exists (see the solid sphere dot), and the free energy is a minimum at *α/π* = 1 for low ionic strengths, e.g., *C*_ion_ = 10 mM. As the ionic strength increases, the metastable states 0 *< α/π <* 1 become the equilibrium configuration. The transition from a global minimum to a local minimum occurs at the threshold value 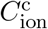. In contrast to ionization, ionic concentration effects show an opposing trend, which is that reducing *C*_ion_ leads to a higher membrane curvature morphology (see the membrane shape contour in Fig. 5B). The sequence of equilibrium membrane morphologies for a sequence of *C*_ion_ values is displayed on the right column in Fig. 5B.

Next, to intuitively visualize how the global minimum energy state is shifted by increasing *ϕ*_b_ and *C*_ion_, we focus on the stable state corresponding to the optimal opening angle *α*_min_. Figures 5C and 5D show the dependence of the equilibrium opening angles *α*_min_ on *ϕ*_b_ and *C*_ion_. Examining the effects of varying the degree of ionization and ionic strength, we note that the optimum opening angle *α*_min_*/π* first shifts gradually, then jumps to unity at the critical values 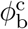 and 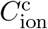. This behavior implies a smooth transition first and then followed by a snap-through transition. Such a sharp jump indicates that increasing the degree of ionization or reducing ionic strength to a specific threshold can trigger the transition of membrane geometry from a spherical cap (open star) to a complete sphere (closed star). Moreover, the degree of ionization and ionic strength also affect the critical grafting density that triggers the snap-through transition, as shown in Fig. S10. Figure 5E maps the equilibrium opening angle in the parameter space of *ϕ*_b_ and *C*_ion_. As a result, all the features of Figs. 5C and 5D can be captured by Fig. 5E. Additionally, the heatmap shows that two regimes corresponding to spherical cap (regime I) and complete sphere (regime II) are identified, in which the solid curves that separate regime I from regime II represent the critical values. The critical line in Fig. 5E shows a proportional dependence relationship between *ϕ*_b_ and *C*_ion_, revealing that reducing the ionic strength lowers the threshold *ϕ*_b_ required to trigger the discontinuous transition in membrane structures. Together, these observations predict that conditions of high degree of ionization and low ionic concentration result in amplified membrane bending and give rise to the formation of higher membrane curvature structures. Collectively, these results suggest that, beyond polymer grafting density and length, changes in ionization and ionic strength provide additional means to modulate the curvature-generating activity of membrane-tethered polymers, enabling cells to tune membrane curvature in response to their ionic environment.

### Experiments reveal pH- and ionic strength-dependent curvature generation

We next examined how environmental electrostatic dependencies influence GUV membrane curvature generation. Consistent with our initial observations, exposing GUVs to 1 *µ*M his-AP180CTD induced the formation of protein-coated membrane tubules protruding outward from the vesicle surface (Fig. 6A, C). Increasing the buffer NaCl concentration from 50 mM to 500 mM substantially reduced the frequency of outward membrane tubulation from 60.6% ± 8.0% to 39.2% ± 4.6% (Fig. 6B, *P*=0.016), consistent with electrostatic screening that attenuates intermolecular repulsion among membrane-bound proteins (*14, 44*). Conversely, increasing the buffer pH from 6.4 to 8.4 yielded a progressive increase in tubulation frequency from 49.1% ± 5.8% to 62.8% ± 5.9% (Fig. 6D, *P*=0.035). Given that AP180CTD has an isoelectric point (pI) of 4.17, this enhancement is likely driven by an increased net negative charge at higher pH, which intensifies lateral electrostatic repulsion between adjacent proteins. A similar sensitivity to ionic strength and pH was observed for the truncated his-AP180CTD-1/3 construct (SI Appendix, Fig. S9). These findings further validate the ionic strength dependencies predicted by our theoretical model.

**Figure 6:**
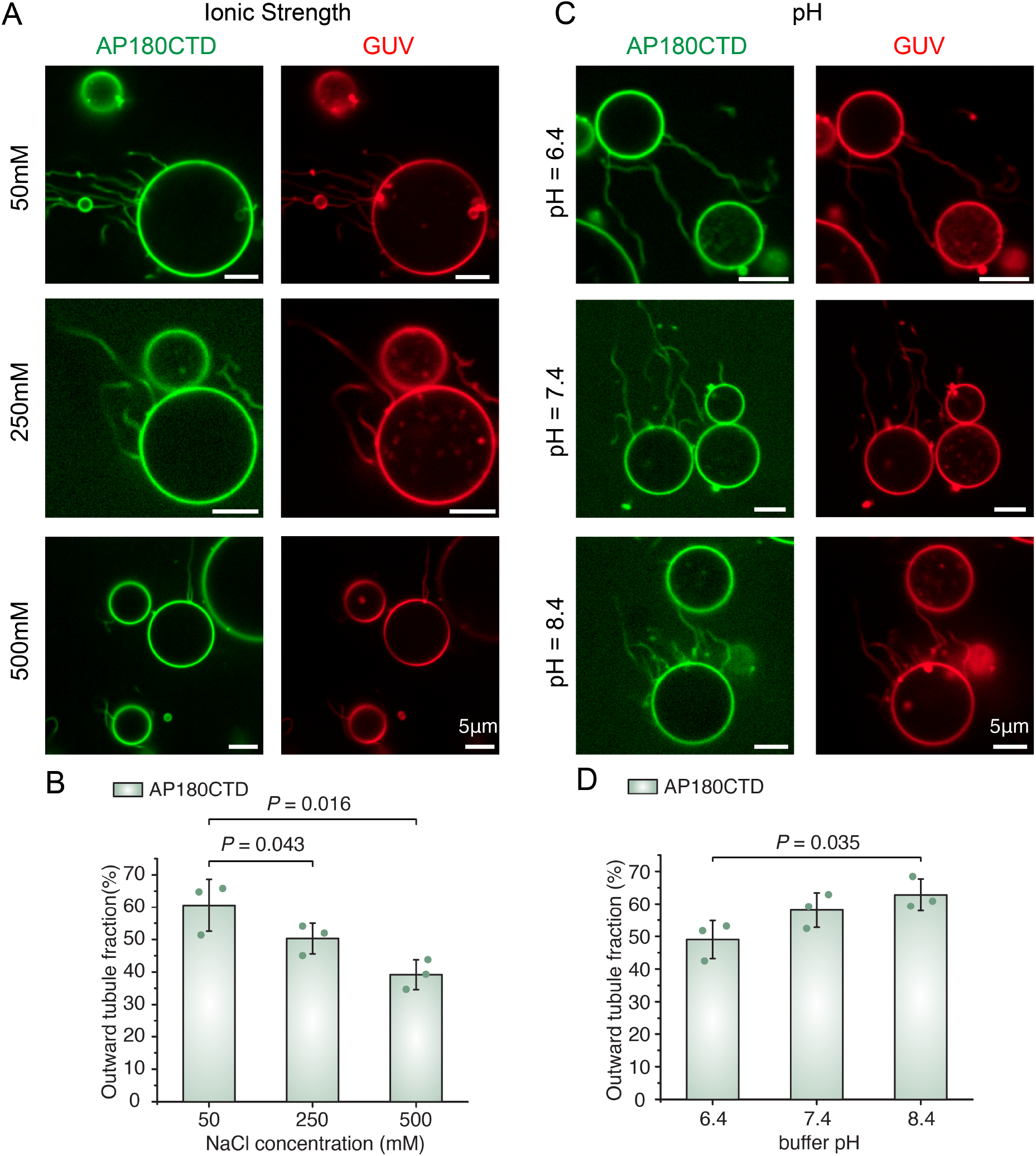
Tubulation of GUVs driven by his-AP180CTD is modulated by ionic strength and pH. (A, C) Representative confocal images (protein and lipid channels) of protein-coated tubules emanating from GUV surfaces. GUVs were incubated with 1 *µ*M his-AP180CTD varying NaCl concentrations (50 mM, 250 mM, and 500 mM) in (A), and varying pH levels (6.4, 7.4, and 8.4) in (C). Scale bar = 5 *µ*m. (B, D) Quantification of the fraction of GUVs displaying outward tubules across the different NaCl (B) and pH (D) conditions. GUV membrane composition was 83 mol% POPC, 15 mol% DGS-NTA-Ni, 2 mol% DP-EG10 biotin, and 0.1 mol% Texas Red–DHPE. Experiments evaluating NaCl variation were conducted in 25 mM HEPES at pH 7.4, while experiments evaluating pH variation were conducted in 25 mM HEPES containing 150 mM NaCl. Error bars represent the standard deviation (SD) of three independent trials (individual data points shown). Statistical significance was evaluated using an unpaired, two-tailed Student’s t-test.

### The height of the polymer layer is modulated by the solution environment

Inferring and measuring the height of densely coated macromolecules on cell membranes can provide useful information for the architectural details and physical properties of the brush-like macromolecular layer. For instance, on the surface of endothelial cells, the height of polymer brush defines the thickness of the glycocalyx. On the inside of the cell, the height of the membrane-tethered polymer layer defines the search radius for endocytic IDRs looking for binding partners. Therefore, we finally asked how the ionic strength and grafting density impact the height of the polymer brush layer *L*_brush_. For this calculation, we leveraged the relationship between brush height and membrane curvature. Briefly, the equilibrium membrane geometry can be determined by identifying the optimal opening angle, *α*_min_, corresponding to the stable state on the total energy landscape. This optimal opening angle determines the curvature of the polymer-coated membrane through 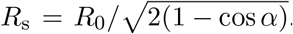. The resulting membrane curvature, 1*/R*_s_, can then be used to predict the brush thickness according to Eq. (S39). We plot the brush height as a function of ionic strength (Fig. 7A) and grafting density (Fig. 7C). Two key observations emerge from the results. First, increasing ionic strength leads to a pronounced reduction of polymer brush height (Fig. 7A). When *ϕ*_b_ = 0.1, the dependence of brush height on ionic strength exhibits two distinct regimes and a non-monotonic behavior, arising from a snap-through transition of the membrane shape from a sphere to a spherical cap, accompanied by a discontinuous change in *R*_s_ (see the inset of Fig. 7A). This transition is similar to that shown in Fig. 5D, where discontinuous changes in *α*_min_ occur. To the left of the dotted blue line, the system behaves like a membrane-tethered polymer brush layer on a sphere of constant radius (Fig. 7B), where a monotonic decrease in brush height is observed with increasing ionic strength. However, when crossing the dotted line, the brush height jumps abruptly to a higher value as a result of the membrane shape transition, corresponding to a first-order transition. For *ϕ*_b_ = 0.2, the orange curve displays a continuous decrease in brush height with increasing ionic strength, for which the equilibrium membrane shape remains a sphere with a constant radius throughout the entire range of ionic strength considered. In solution with low ionic strength, the height of the polymer brush is extended considerably, owing to the magnitude of the exclude volume interactions being enhanced at large electrostatic effect inside the brush layer, causing the polymer chain to expand. However, in high ionic strength conditions, this electrostatic effect is screened, effectively reducing the exclude volume interactions and, therefore, resulting in a shrinkage of the polymer brush thickness. Notably, when the ion concentration becomes sufficiently high, further increases in ionic strength appear to have little effect on brush height. As demonstrated in Fig. S4, variations in ionic strength and the degree of ionization effectively modulate the excluded-volume interactions between polymer chains in the brush layer. Therefore, our results further suggest that, for negatively charged membrane-tethered polymers, the electrostatic effect generated by the negative charges in the polymer brushes serves to enhance the steric effects.

**Figure 7:**
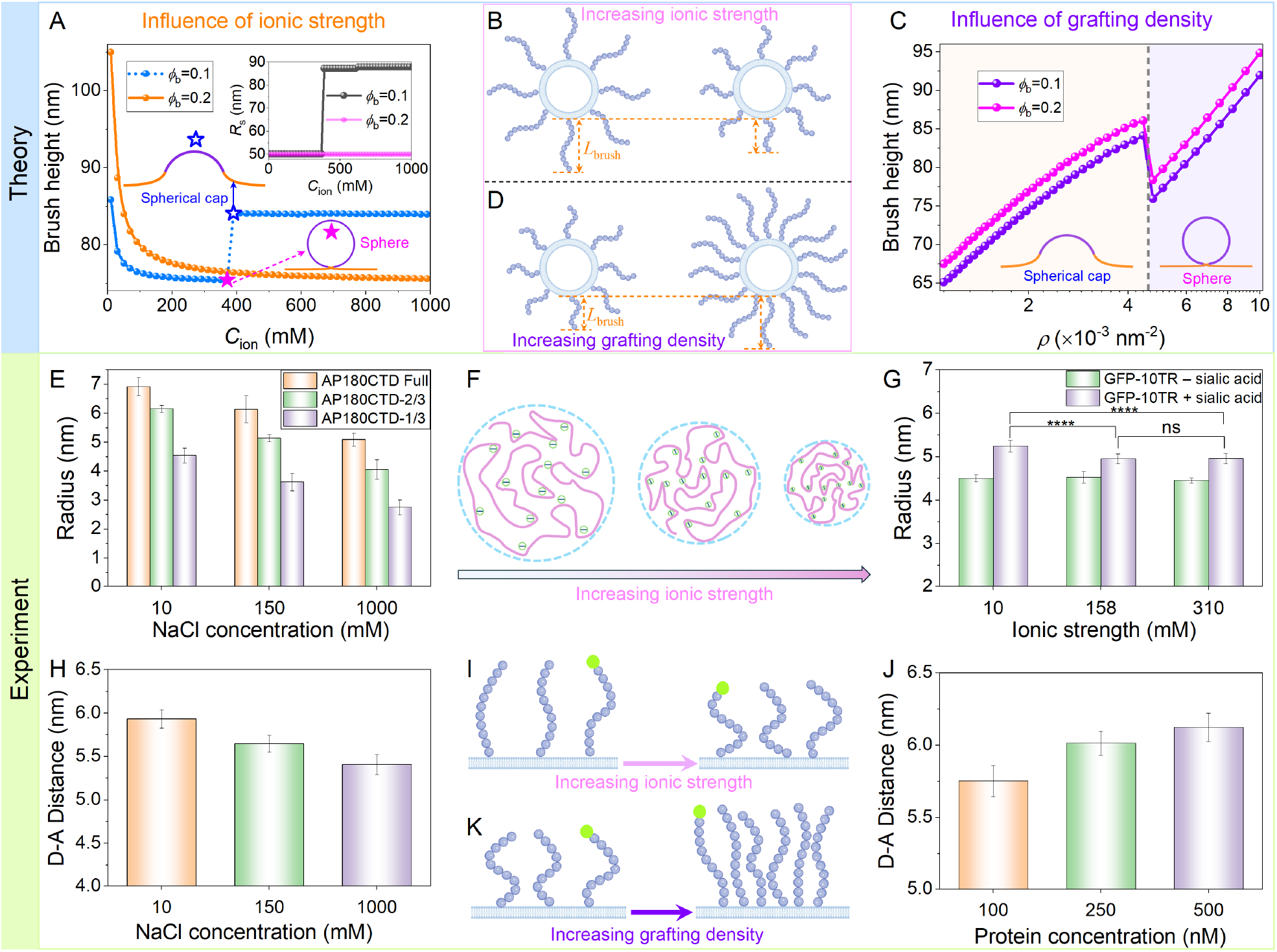
Membrane-tethered polymer brush height on a membrane surface. Brush height as a function of (A) ionic strength *C*_ion_ when *ρ* = 4.76 *×* 10^−3^ nm^−2^, and (C) grafting density *ρ* at *C*_ion_ = 150 mM, for two different degrees of ionization, *ϕ*_b_ = 0.1 and *ϕ*_b_ = 0.2. The remaining parameters are listed in Table S1. The inset in (A) shows the dependence of membrane radius on ionic strength. Schematic of the height variation of brush layer on a spherical membrane geometry under (B) different ionic strengths and (D) grafting densities. Hydrodynamic radius of (E) AP180CTD truncation mutants and (G) GFP-10TR protein with and without sialic acid at various salt concentrations. Experimental data in (E) and (G) are cited from Ref. (*14*) and Ref. (*48*), respectively. (F) A schematic representation of the impact of ionic strength on polymer chain hydrodynamic radius. Donor–acceptor distance as a function of (H) NaCl concentration and (J) protein concentration. Experimental data in (H) and (J) are replotted from Ref. (*19*). Schematic of intrinsically disordered protein deforms in response to changes in (I) ionic strength and (K) crowding.

Analyzing the dependence of brush height on grafting density, we find that brush height increases with increasing grafting density, as expected (Figs. 7C and 7D). For densely crowded membrane-tethered polymers, steric repulsive interactions between the chains drive the extension of the polymer brush away from the membrane surface. This result is qualitatively in good agreement with both experimental observations and molecular dynamics simulations reported in (*24*), where the average height of a multidomain protein was measured at three different surface densities. Again, a discontinuous jump in brush height occurs at the dashed line, which separates two distinct regimes: the left-hand regime corresponds to spherical cap membrane shapes, whereas the right-hand regime corresponds to spherical membrane shapes. In analogy to Fig. 3C, this transition is accompanied by discontinuous changes in *α*_min_ and *R*_s_, as shown in Fig. S11. Following this shape transition, each polymer chain gains more available space on the more highly curved spherical membrane surface, reducing the steric crowding between polymer chains and thereby giving rise to an immediate decrease in brush height after the transition (Fig. 7C). A direct comparison of brush heights at different degrees of ionization reveals that higher ionization tends to promote membrane-tethered polymer extension (Fig. 7C). Together, we predict that increasing ionic strength reduces the brush thickness, whereas higher grafting density and ionization increase it.

Next we sought to compare these predictions to available experimental data. First, to gain a better understanding of the impact of ionic strength on polymer hydrodynamic radius, independent of the membrane surface, we examined measurements of the hydrodynamic radius of the AP180CTD truncation mutants and GFP-10TR, free in solution, with and without sialic acid at different salt concentrations. For intrinsically disordered proteins, Fig. 7E shows that the hydrodynamic radius of each AP180CTD truncation mutant decreases with increasing sodium chloride concentration, indicating that interchain electrostatic interactions can be modulated by varying the ionic strength of the solution (Fig. 7F). Similarly, for tandem repeats (TRs) of Mucin without sialic acid, the hydrodynamic radius does not change significantly as the salt concentration increased from 10 mM to 310 mM (Fig. 7G). In contrast, the hydrodynamic radius of the fully glycosylated GFP-10TR ligand containing sialic acid decreases with increasing ionic strength from 10 mM to 158 mM (Fig. 7G). Together, these observations demonstrate that the physical properties of charged biopolymers are significantly impacted by electrostatic interactions. Subsequently, for polymers grafted onto the membrane surface, we examined measurement of polymer extension as a function of ionic strength and protein concentration. Previous work empolyed a FRET (Förster resonance energy transfer)-based sensor to measure the distance between the free and membrane-bound ends of tethered polymers (*19*). This sensor consisted of a PEG (polyethylene glycol) chain that was covalently attached at one end to the headgroup of a synthetic phospholipid. The free end of the PEG chain was labeled by a donor fluorophore, while lipids labeled with acceptor fluorophores resided at the membrane surface. Using this system, the characteristic donor-acceptor distance between the two ends of the disordered protein chain at different ionic strengths and protein crowding conditions was estimated, as shown in Figs. 7H and 7J. Our theoretical prediction is consistent with these experimental results reported by Houser et al. (*19*), for the extension of anchored proteins on membranes, as a function of ionic strength (Fig. 7H) and protein concentration (Fig. 7J). These experimental results show that the donor–acceptor distance decreases when ionic strength of the solution is increased from 10 mM NaCl to 1 M NaCl (Figs. 7H and 7I), and increases when protein concentration is raised from 100 nM to 500 nM (Figs. 7J and 7K), a conclusion that aligns with our findings. Taken together, these experimental results strongly support the predictions of our theoretical model.

## Discussion

In this study, we developed a unifying model that is able to explain the ability of diverse membrane-tethered polymers to sense and generate membrane curvature in response to changes in solution and surface crowding conditions. Our comprehensive framework contributes to a physical understanding of how membrane-tethered polymers generate steric repulsion at membrane surface. For densely crowded membrane-tethered polymers, a decrease in ionic strength or increase in grafting density or the degree of ionization enhances the collision and crowding of polymer chains on membrane surfaces. The consequently stronger steric repulsions between polymer chains in the crowded environment can stretch the polymer and bend the membrane, and lead to the distribution of membrane-tethered polymers in higher membrane curvature regions. Our model predicts that increasing polymer length enhances the partitioning number, while increasing ionic strength has the opposite effect. This prediction indicates that membrane-tethered polymers prefer regions of higher local membrane curvature. Examining the thickness of the brush layer, we find that the height of membrane-tethered polymers strongly depends on ionic strength, the degree of ionization, and the polymer grafting density. Predictions from our model are qualitatively consistent with a large number of experimental observations beyond those used in this study. For example, our model predictions are applicable to the study by Bracha et al. (*49*), who measured the height of dense DNA polymer brushes on a biochip using evanescent fluorescence and found that the brush compresses with increasing ionic strength and extends with increasing grafting density. Our findings also align with previous predictions that charged multidomain proteins swell relative to their uncharged counterparts (*25*). In addition, the height of the thick glycocalyx of epithelial or endothelial cells has previously been estimated to be approximately 100 to 500 nm (*50, 51*). Here, the height (~ 100 nm) of membrane-tethered polymers evaluated by the present model agrees, in terms of order of magnitude, with those measured in experiments (*50, 51*). Thus, our model is valid for a wide range of experimental observations.

Curvature sensing of membrane-tethered polymers can be understood as an entropic mechanism in which highly curved membranes provide more available space and greater orientational degrees of freedom for polymer chains, allowing membrane-tethered polymers to maximize their conformational entropy by assembling on membrane surfaces with higher curvature (*17*). Therefore, the agreement between our predictions and experimental observations (Fig. 2) suggests that our theoretical model can be used to explain curvature sensing for many membrane-tethered polymers (e.g., mucins (*15*)) over a wide range of ionic strengths and polymer lengths. For curvature generation, the prevailing view has been that membrane bending is driven by the following three main mechanisms: wedging, scaffolding, and crowding. In contrast, here we show that membrane-tethered polymers forming a brush-like network on membranes are also potent drivers of bending. The layer of polymer brush-like structures exerts an entropic bending force on the membrane surface, generating a diverse family of membrane shapes. This behavior arises because, on a membrane with a high grafting density of polymer brush networks, collisions between adjacent polymer chains stretch the chains perpendicularly to the membrane surface (*4, 26, 52*). This stretching reduces chain entropy and consequently increases the chain free energy (*4, 26, 52*). To reduce the entropy cost, the membrane bends toward the polymer side, thereby creating more space for the polymer chains and alleviating their stretching. As a result, a repulsive interaction between the chains (or an effective entropic bending force) is exerted on the membrane, leading to curvature driven by the accumulated entropic free energy from stretching the membrane-grafted polymer brushes. From this perspective, membrane-tethered polymers can be viewed as crowding agents with a propensity to occupy larger volumes compared to many other membrane curvature generators, such as scaffolding proteins. Consequently, the curvature generated by membrane-tethered polymers is influenced by multiple factors, including polymer grafting density, polymer length, degree of ionization, pH, and ionic strength. Notably, the impact of degree of ionization, pH, and ionic strength on curvature generation has not been incorporated in previous studies (*4, 16, 26*–*36*). Modulating these factors provides a means to control the curvature-generating ability of membrane-tethered polymers, thereby lowering the total free energy of the membrane-polymer composite. This suggests a possible mechanism by which cells endowed with dense membrane-tethered polymers may tune their curvature-generating activity in response to changes in ionization and ionic strength. Based on these observations, we propose a mechanism to explain the role of ionic strength: increasing ionic strength amplifies the electrostatic screening effect, which weakens the effective excluded volume interaction and consequently leads to compression of the membrane-tethered polymer layer.

Importantly, our findings have implications for various membrane trafficking processes, particularly those involving outward membrane bending (away from the cytosol), such as microvesicle formation, filopodia formation, and cellular blebbing (*4, 53*–*55*). Although the roles of cortex attachment, cytoskeletal assembly, and filament bundles have been investigated in these studies, the impact of membrane-tethered polymers remains to be elucidated (*55*–*59*). It is possible that membrane-tethered polymers inherently provide an effective entropic bending force that drives outward bending. It is also possible that the favorable energetic contribution of membrane-tethered polymers to outward bending can overcome the actin-membrane cortex attachment when blebs or extracellular vesicles need to form (*36*). For example, in experiments (*4*), cryo-TEM confirmed that Muc1-42TR ΔCT-expressing cells secrete glycocalyx-decorated membrane vesicles, and confocal imaging showed that functionalizing GUVs with Podxl’s S/T-rich domain induces similar spherical structures. This might have implications for understanding microvesicle generation in many cancer cell types, such as tumor cells with high mucin expression tend to generate a large number of microvesicles (*60*). Additionally, in vitro experiments exhibit evidence that the steric pressure generated by intrinsically disordered proteins can drive vesiculation and breakup of membrane vesicles to form smaller daughter vesicles (*19, 61*). Together, these studies indicate that, using this steric entropic mechanism, membrane-tethered polymers on membrane surfaces contribute to outward membrane bending. Moreover, the predicted dependence of curvature generation on various system parameters may provide guidance on how to adjust control parameters to fine-tune membrane geometries in experiments.

Finally, our model provides a framework not just for understanding curvature sensing and generation, but also for the extension of membrane-tethered polymers on membrane surface. Although the effects of ionic strength and ionization on these processes have been observed experimentally, a model capable of predicting their effect has been lacking. In particular, both our theoretical model and experiments suggest that curvature sensing and generation, and brush extension abilities are improved upon elevation of ionization degree and reduction of salt concentration, which we attribute to the enhancement of effective excluded volume interactions. We note that membrane-tethered polymer interactions can also lead to inward membrane bending under some conditions (*2, 44, 62*). Therefore, future modeling and experimental studies need to incorporate additional biological details of the cellular plasma membrane, including the spatial heterogeneity of membrane-tethered polymers distribution, to dissect the contributions of different molecular moieties to the direction of curvature. Our approach has certain limitations and simplifications, such as prescribing the shape of the membrane with polymers grafted as a simplified geometrical scheme and neglecting the spatial heterogeneity of the distribution of the membrane-tethered polymers on the membrane. Recently, tools like proximity biotinylation and cross-linking mass spectrometry have been able to provide data on the type and even the local density of proteins within curved membrane structures. With these data in hand, it would be possible to extend our model to determine where and to what extent biopolymer crowding contributes to membrane curvature generation and sensing in specific biological contexts.

We also acknowledge that the membrane geometries considered in the model do not exactly match those probed experimentally. Although these geometries are not quantitatively equivalent, they capture the same underlying coupling between membrane curvature and polymer steric interactions, which can explain the qualitative agreement between theory and experiment. The model presented here provides a useful guide for understanding how membrane-tethered polymers can sense and generate curvature under different conditions because we were able to bring together disparate experimental findings that have used different proteins and assays under one common theoretical framework. As a result, we can show that they can all be explained by a single set of relatively simple physical assumptions about how polymers interact on a flexible membrane surface. More broadly, this framework offers a physical basis for interpreting and predicting how electrostatic and steric interactions among polymers on membranes tune curvature sensing and generation, and it provides a foundation for future experiments and models that can further resolve geometry-specific effects.

## Data, Materials, and Software Availability

All study data are included in the article and/or SI Appendix. The code is available on https://github.com/RangamaniLabUCSD/membrane-tethered_polymers_ionic_strength.

## Acknowledgments

This work was supported by NIH NIGMS R35GM158446 and NSF 2327423 to P.R. and J.C.S was supported by R35GM139531, National Science Foundation 2529782 and 2327244 to J.C.S., and the Welch Foundation F-2257 to J.C.S. Dr. Yuan is a Robertson Foundation Fellow of the Life Sciences Research Foundation. W. F. Z. was supported NIH/NIGMS R35GM147333.

## Author contributions

K.X. and P.R. conceived the theoretical framework and derived the governing equations. K.X. conducted the simulations and made the figures. W.Z., J.H. Y.F, and A.K. conducted or reanalyzed the experiments. K.X., P.R, Y.F., J.C.S. wrote the paper and all authors edited the manuscript. P.R. and J.C.S. oversaw the theoretical and experimental portions respectively and were responsible for funding acquisition.

## Supporting Information Text (SI)

### 1 Model for ionic strength-dependent curvature sensing of biopolymers

The curvature-sensitive localization of proteins on membranes is essential for the initiation and assembly of curved membrane morphologies and thus is vital for many cellular processes (*63*). For example, assembly of endocytic pit, viral bud, and tubular organelles requires that proteins are capable of sensing and stabilizing membrane curvature (*64, 65*). Curvature sensing refers to the preferential binding or partitioning of proteins to regions of high membrane curvature. To date, many proteins with specific structural features, such as dynamin protein (*41, 66*), epsin protein (*67, 68*), and crescent-shaped Bin Amphiphysin Rvs (BAR) protein (*69*–*71*), have been identified as sensors and generators of membrane curvature. Recently, intrinsically disordered proteins, which lack a defined three-dimensional fold, and glycoproteins such as MUC1 have been also discovered to exhibit curvature-sensing behavior (*14*–*16*). These disordered proteins behave like random polymers, and the glycoproteins are in a polymer brush network. In addition, the ionic strength plays an important role in cellular physiology. For instance, ionic strength influences a wide array of cell biological processes, such as determining the structure of intrinsic disordered proteins (*72*), regulating the activity of enzymes (*73*), affecting protein aggregation and phase separations (*74, 75*), and associating to membrane curvature generation (*19, 44*). However, the role of ionic strength in the curvature sensing capability of biopolymers and disordered proteins is much less understood. Therefore, understanding and quantifying the effect of ionic strength on the curvature sensing of biopolymers is of crucial importance to many cellular processes under physiological conditions.

#### 1.1 Thermodynamic model of biopolymer Binding to membrane tubes

To investigate the effect of ionic strength on the curvature sensing of biopolymers, we present here a thermodynamic model to describe the relative partitioning of the attachment of biopolymer to membrane tubes and to the cell surface. To derive the thermodynamic model, we first need to introduce the free energy of the membrane in the presence of bound biopolymers for a curved membrane tube (e.g., filopodia) connected to a cell body (e.g., a giant unilamellar vesicle (GUV)). We then calculate the enrichment of biopolymers on the tube relative to the vesicle at equilibrium. Finally, our model shows that the variation trend of relative partitioning as a function of ionic strength is in agreement with experiments.

##### 1.1.1 Free energy of membrane with adsorbed biopolymer

The free energy of the membrane in the presence of biopolymers is composed of the sum of six terms, i.e.,

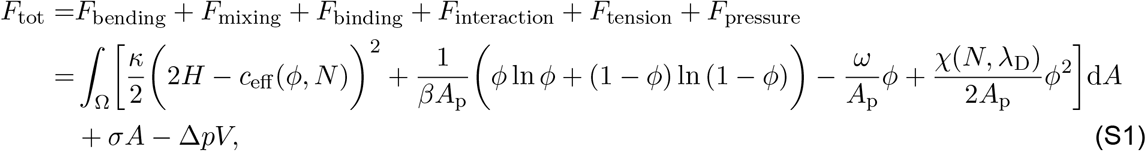

where *c*_eff_ is the effective spontaneous curvature of the membrane–generated due to the steric interaction of the biopolymer, *A*_p_ is the surface area occupied by the biopolymer anchor, *ϕ* is the area fraction of biopolymer on the cell membrane, *ω* represents the chemical potential of the biopolymer attachment to the membrane, and *χ* is a direct interaction coefficient between biopolymers, Δ*p* is the hydrostatic pressure difference across the cell membrane. Because we have established that the biopolymer can generate curvature, it is reasonable to lump together the curvature generation effects of the biopolymer into an effective spontaneous curvature, *c*_eff_. Since *c*_eff_ is a phenomeno-logical coefficient related to polymer area fraction and its length, we assume that this effective spontaneous curvature is a function of the polymer area fraction *ϕ* (ratio between the membrane area anchored by polymers and the area of the entire membrane) and the number of effective monomeric segments *N*. There are many models in the literature for the curvature induced by different polymer systems on the membrane (*26, 27, 30*–*34, 76*). We chose the model presented in (*76*) because it incorporates the effect of *ϕ* and *N* on the spontaneous curvature

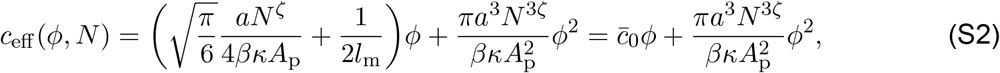

where *l*_m_ is the membrane thickness (*l*_m_ 4 5 nm), and the exponent *ζ*= 3*/*5 for polymer brushes under good solvent conditions. Here, for simplicity, we merely consider the linear term in Eq. (S2) at low polymer densities.

Also, note that our analysis restricts the biopolymer-membrane system immersed in an ionic solution environment. Consequently, this will introduce electrostatic screening, characterized by the Debye length *λ*_D_. Many experimental observations show that a higher ionic strength leads to a smaller hydrodynamic radius, indicating that the Debye screening length, *λ*_D_, is reducing. Thus, a higher ionic strength results in a facilitation of the chance that the monomers encounter each other between inter-polymers as the reduced electrostatic screening effect. As a consequence, the interaction between biopolymers can be changed due to the variation of Debye screening length, *λ*_D_, under different ionic conditions. In this case, to take into account the effect of ionic strength on curvature sensing, we speculate that the Flory interaction parameter *χ* is a function related to the polymer length *N* and the Debye screening length *λ*_D_, i.e., *χ* = *χ*(*N, λ*_D_). However, the exact dependence of *χ* on *N* and *λ*_D_ is still unknown for biopolymers. For simplicity, we assume that they obey a linear dependence given by 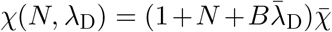, where *B* is a phenomenological coefficient which is assumed to be equal to *N* in our numerical simulations, 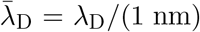 denotes the dimensionless Debye screening length, and 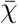 is a measure of the strength of inter-polymer interactions (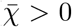 and 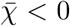 represent repulsive or attractive interactions respectively). Here, the Debye length is dependent on the ionic strength (salt concentration) *c*_s_ of the solution via the following relation

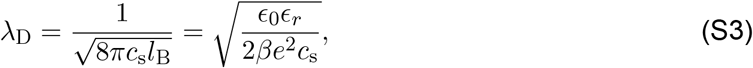

where *l*_B_ = *βe*^2^*/*(4*πϵ* _0_ϵ _*r*_) is the Bjerrum length with *e* being the element charge, _0_ being the vacuum permittivity, and _*r*_ being the relative permittivity. In such a treatment, the coefficient *χ* can be reduced to the frequently encountered case that the effects of biopolymers size and ionic strength are neglected, corresponding to 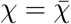 when *N* = 0 and *λ*_D_ is excluded.

The proposed Eq. (S1) also applies to the case when a membrane is attached to the underlying cortex and the membrane tube is pulled by an external force.

##### 1.1.2 Biopolymer partitioning

In Eq. (S1), we write the individual energy contribution in a generic form that can be applied to the membrane tube or to the vesicle. Below, we consider the biopolymer-membrane system consisting of a vesicle component that represents the whole cell and a long membrane tether of radius *R*_t_ and length *L*_t_ connected to the cell body, as shown in Fig. S1. The first energy term on the right-hand side of Eq. (S1) is the membrane bending energy which is given by

**Figure S1:**
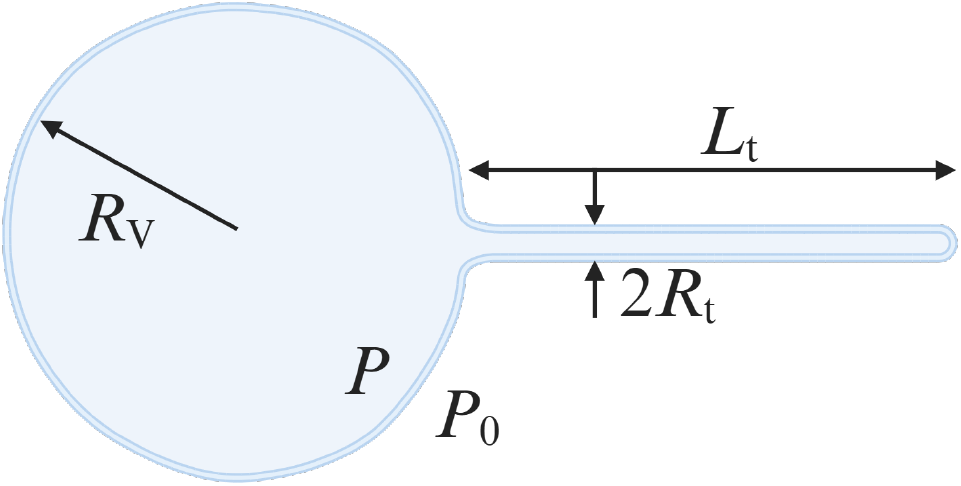
Schematic of the system as used in the curvature sensing model. We consider the membrane to consist of a vesicle component that represents the whole cell and tubular component with radius *R*_t_ and length *L*_t_.

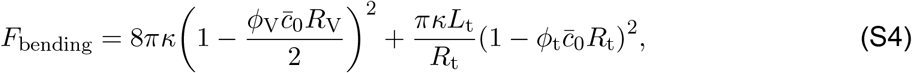

where *ϕ*_V_ and *ϕ*_t_ are the biopolymer area fractions on the sphere cell and tube, and *R*_V_ is the radius of the spherical cell. The second term appearing in Eq. (S1) represents the mixing energy, which has the Flory-Huggins form, as follows

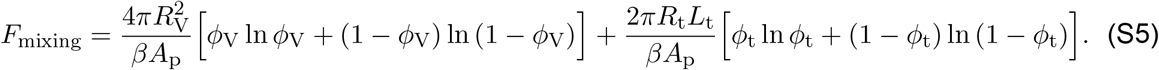

The third term is the energy associated with biopolymer binding to the membrane

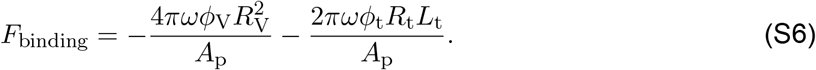

The fourth term takes into account the energy arising from polymer-polymer interactions and is given by

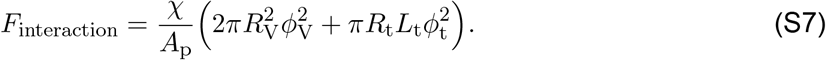

The fifth term represents the surface tension energy and is given by

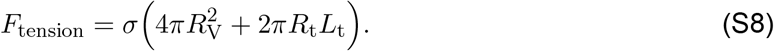

The last term is the work done by the hydrostatic pressure difference, Δ*p*, between the inside and outside of the cell

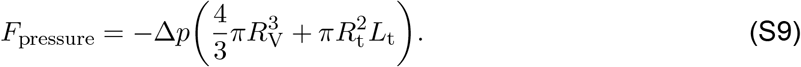

Therefore, summing Eqs. (S4)-(S9) leads to the total energy

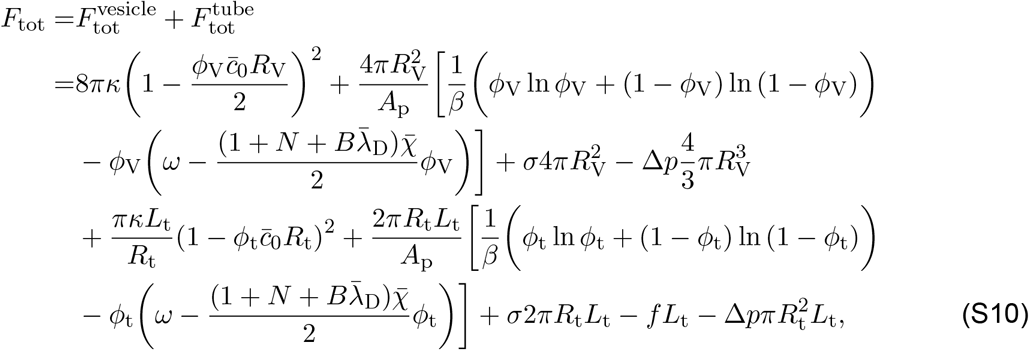

where

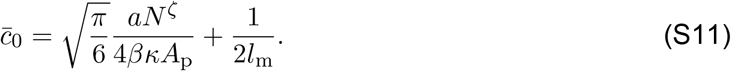

To quantify the curvature-sensing ability of biopolymers, we need to determine the relative partitioning of biopolymers on the tube and on the cell surface, which is defined by the ratio *ϕ*_t_*/ϕ*_V_. In thermal equilibrium, the polymer-bound fraction *ϕ* is locally determined for the given curvatures. By minimizing the total free energy with respect to *ϕ*_V_, i.e., ∂*F*_tot_*/*∂*ϕ*_V_ = 0, one can obtain

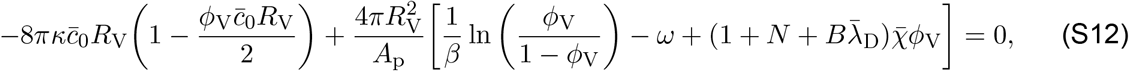

Assuming that the vesicle (or the cell) can be treated as nearly flat, 1*/R*_V_ = 0, the above equation reduces to

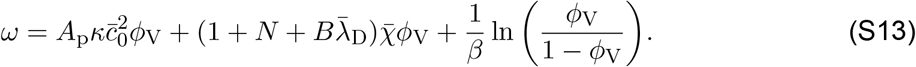

which can be solved numerically to obtain *ϕ*_V_. Minimization of *F*_tot_ with respect to *R*_V_ then will leads to the Laplace law

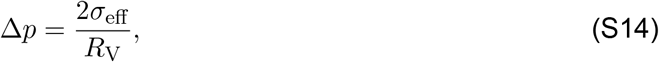

where

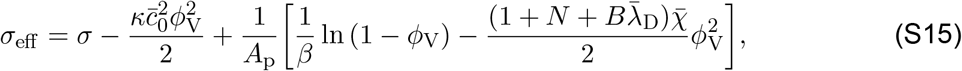

Similarly, the equilibrium fraction of polymers on the tube can be obtained by minimizing *F*_tot_ with respect to *ϕ*_t_, which yields

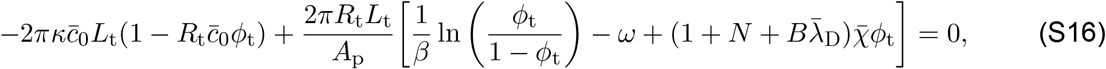

Substituting Eq. (S13) into Eq. (S16) and then expanding it to the first order in Δ*ϕ* = *ϕ*_t_ − *ϕ*_V_, one can obtain

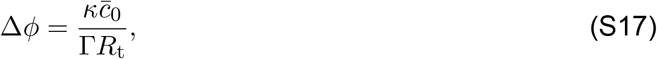

where

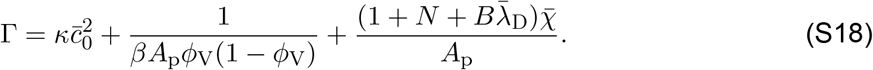

Next, minimizing *F*_tot_ with respect to *R*_t_ and neglecting the pressure term leads to

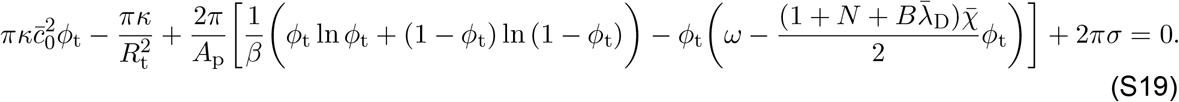

Expanding the above equation to the second order in Δ*ϕ* = *ϕ*_t_ − *ϕ*_V_ and using Eqs. (S13) and (S17) yields the tube radius

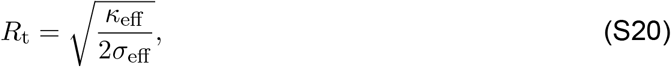

where

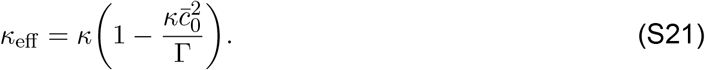

Note that the term, 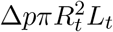, has been neglected due to that it is of order *R*_*t*_*/R*_*V*_ smaller than the other terms (*38*).

Based on these derivations, we minimize the total energy *F*_tot_ with respect to *ϕ*_V_, *R*_V_, *ϕ*_t_, and *R*_t_, respectively, to determine the relative partitioning of biopolymers on the tube and on the cell surface, *ϕ*_t_*/ϕ*_V_. We can obtain *ϕ*_V_ by numerically solving Eq. (S13), and then determining *ϕ*_t_*/ϕ*_V_ = 1 + Δ*ϕ/ϕ*_V_ by using Eq. (S17). This allows us to investigate the dependence of *ϕ*_t_*/ϕ*_V_ on ionic strength (salt concentration) *c*_s_ and biopolymer length *N*, as shown in Figs. 2B and 2C in the main text.

### 2 Elastic and line tension energy contributions of the membrane with biopolymers grafted

#### 2.1 Elastic energy of the membrane patch grafted with glycocalyx

Specifically, we treat the cell membrane as a homogeneous thin elastic layer. To describe the energy cost of bending the membrane away from its preferred shape, we use the well-known Canham-Helfrich energy density (*42*), *f*_bending_ = *κ*(2*H*)^2^*/*2, where *κ* is the membrane bending rigidity, *H* = (*c*_1_ + *c*_2_)*/*2 is the mean curvature in which *c*_1_ and *c*_2_ are the two principal curvatures. For the membrane patch grafted with glycocalyx, since the membrane shape of this part is restricted to a spherical cap geometry, the two principal curvatures are given by *c*_1_ = *c*_2_ = 1*/R*_*s*_, the excess area is calculated as 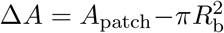, where 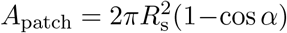 is the area of the spherical cap and *R*_b_ = *R*_s_ sin *α* is the in-plane base radius, and the length of the domain boundary can be written as ∮^∂*l*^ d*l* = 2*πR*_b_. Therefore, the bending energy of the membrane patch with polymer-grafted is calculated as

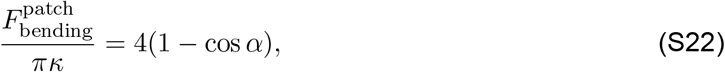

where dividing by *πκ* normalizes the energy.

Remodeling the membrane away from an initially flat shape also involves a surface energy contribution, *F*_tension_ = *σ*Δ*A*, which corresponds to the work required to form a curved membrane shape against a lateral membrane tension *σ*, where Δ*A* denotes the excess area compared to a flat membrane. Thus, the contribution of the tension to the energy of this part is

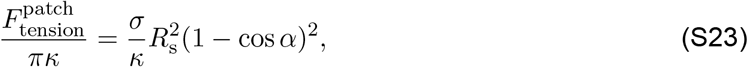

Summing Eq. (S22) and Eq. (S23) leads to the elastic energy of the membrane patch grafted with biopolymers

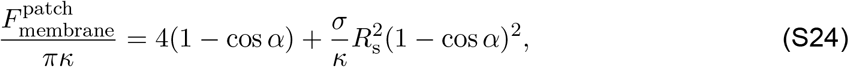

which corresponds to the first two terms in Eq. (3) in the main text.

#### 2.2 Line tension energy

To take into account the multiple phases on the membrane, for example, the polymer rich domain and the surrounding bare membrane, we also considered the effect of the line tension at the biopolymers anchored domain boundary or interface ∂*l*. Hence, the interfacial energy or the energy associated with the line tension is given by *F*_line tension_ = *λ ∮*^∂*l*^ d*l*, where *λ* denotes the strength of line tension along the domain boundary between the biopolymer attached and unattached region, and the integral is over the periphery line d*l* of the domain. We assume that the line tension is a constant. Based on the prescribed membrane geometry, the line tension energy is given by

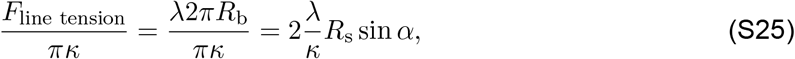

which corresponds to the third term in Eq. (3) in the main text.

### 3 Derivation of the energy contribution associated with the biopolymers

In this section, we briefly recapitulate the derivation of the energy contribution associated with the biopolymers. The reader can also refer to Refs. (*16, 36, 77*) for more details. It is worth noting that our derivation here incorporates the ionic energy contribution, which was not taken into account in the previous model (*16, 36*).

Here we provide a detailed derivation of the fourth term of Eq. (3) presented in the main text and how to obtain the shape equations via the variational methods. Notably, the crowding of large glycosylated proteins appears to regulate the shape of the underlying bilayer plasma membrane (*4*). According to polymer physics, two regimes can be defined depending on the grafting density of biopolymers on the cell membrane, which are the mushroom-like regime and the brush-like regime. Shurer et al. (*4*) have reported that the mucins are in the coiled mushroom state in the case of low-density biopolymers, i.e., glycocalyx. We focus on densely grafted regions of the membrane and therefore model the biopolymers in the brush regime.

**Figure S2:**
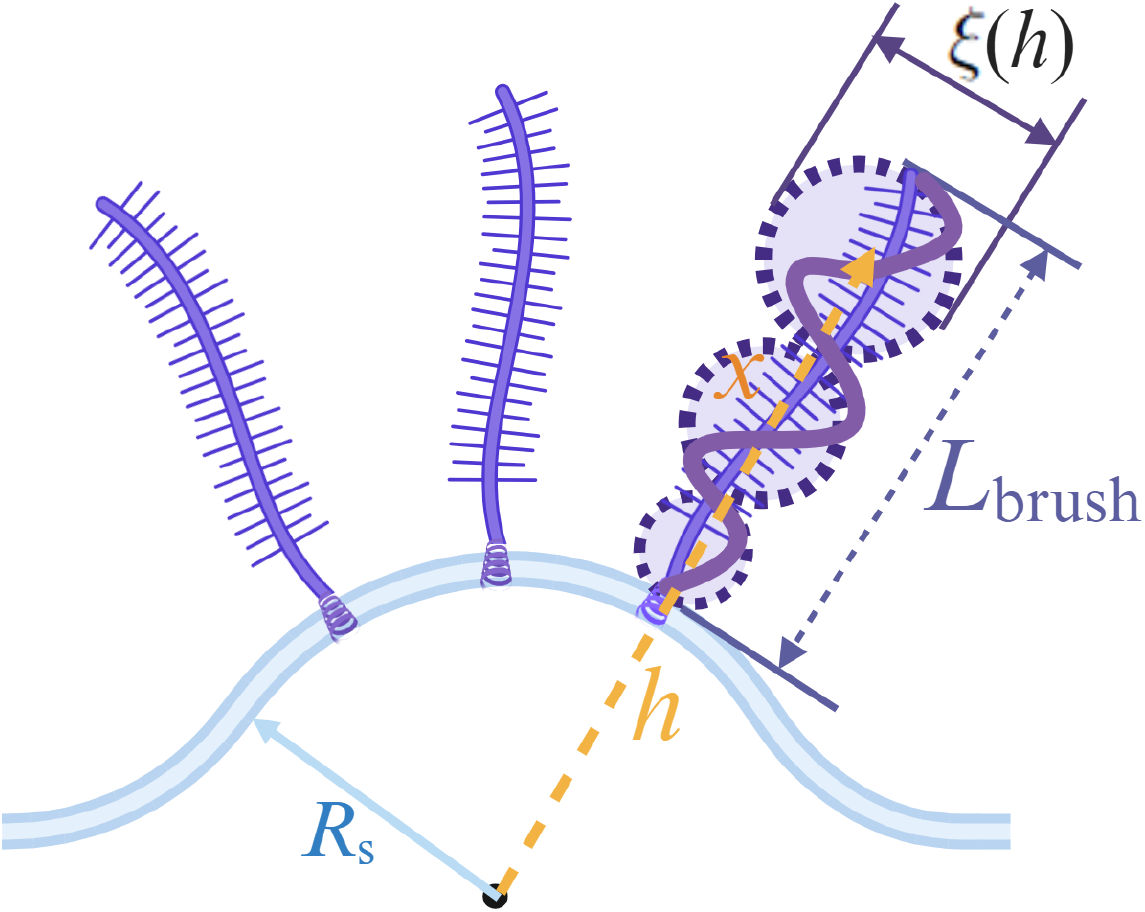
Schematic of a layer of brush-like polymers attached to a spherical cap membrane surface with radius *R*_s_. At a given position *h*, the coarse grained blob (dashed black circle) size in a polymer brush is *ξ*(*h*), the thickness of the brush is *L*_brush_.

Here, we consider a layer of biopolymers grafted on a membrane of radius *R*_s_ in a brush-like structure with thickness *L*_brush_, as shown in Fig. S2. From the viewpoint of coarse graining, the polymer brush is envisioned as an array of blobs. The size of each blob, *ξ*, at a given position *h* equals the square root of the local area per chain *S* (*h*), where *h* = *x* + *R*_s_ is the radial distance which is defined from the center of the spherical surface, and in which *x* is the distance from the membrane surface. Thus, the blob size 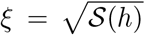 grows as a function of *h*, and the grafting density of polymer brush on the membrane surface can be obtained as *ρ* = 1*/ξ*^2^. Assuming that the layer of biopolymers is of the same height, then the area per chain at distance *x* from the spherical shaped membrane surface is given by (*77*)

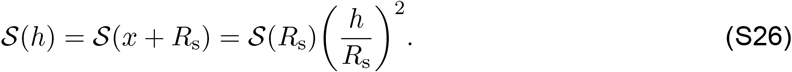

When the membrane is bent, the changes of polymer configuration give rise to the local extension of the polymer chain. Here the local chain extension at a height *h* is characterized by d*h/*d*n*, where the variable *n* denotes the current monomer. This local extension is related to local density profile of monomers *c*_p_(*h*) as (*77*)

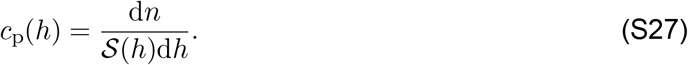

Then, the thickness of the brush, *L*_brush_, is found from the conservation condition (the constraint of conservation of the total number of monomeric segments *N*) (*77*),

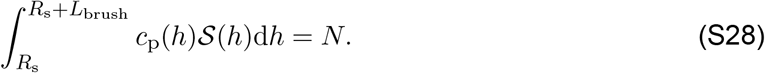

As a result, in the brush regime, the energy contribution originating from biopolymers includes three terms (*4, 77*): the elastic energy of the polymer chain 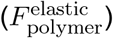, the free energy caused by the excluded volume interactions of polymer monomers 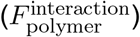, and the electrostatic contribution for the ionic brush 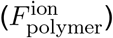. Based on the hypotheses in the main text, according to Ref. (*77*), the elastic energy per chain in the brush can be represented as

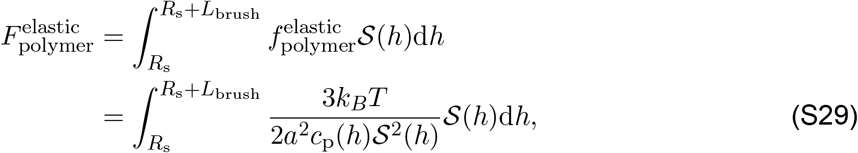

where *a* is the average effective monomeric segment size and 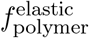 is the elastic energy density of the polymer chain. Within the mean-field approximation, the energy density of the excluded volume interactions (van der Waals interactions) between monomers can be modeled in terms of the virial expansion

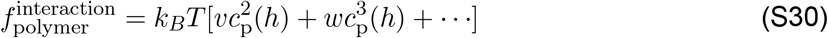

where *v* and *w* are the second and third virial coefficient, respectively. We assume *v* = *a*^3^ *>* 0, implying that the monomer–monomer interactions are purely repulsive. As a result, the excluded volume interactions between monomers per chain in the brush is given by

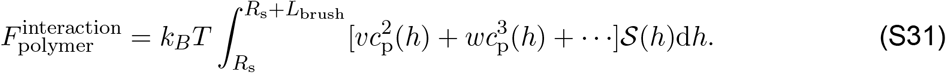

By combining the mean field and the local electroneutrality approximations (*78*), the ionic contribution to the free energy is given by (*4, 44, 77, 78*)

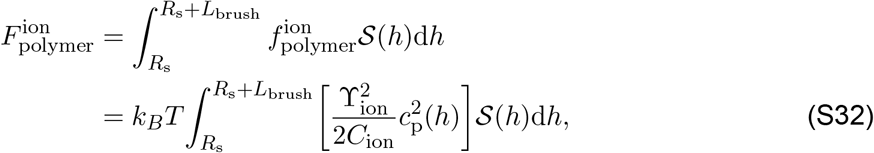

where ϒ_ion_ = *ϕ*_b_ *·N*_ion_ is the degree of ionization of each segment, and 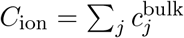 is the ion concentration (ionic strength) in the bulk solution, which is multiplied by the Avogadro’s constant *N*_A_ in the simulations. Here, *ϕ*_b_ is the degree of ionization of each amino acid in biopolymers, and *N*_ion_ = *a/a*_m_ denotes the number of ionizable sites per segment with *a*_m_ being the size of a monomer in the polymer chain. Note that the degree of ionization of an acid group in the bulk solution, *ϕ*_b_, is determined by the bulk concentration of protons through the form (*79, 80*)

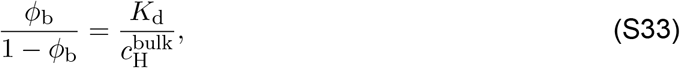

where *K*_d_ is the dissociation constant of acid group within the polymer chain and 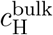 is the bulk concentration of protons, which is related to the pH value of the buffer solution through the form 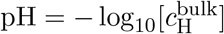. For simplicity, we ignore the possible dependence of *K*_d_ on other parameters and assume that *K*_d_ is a constant. The dimensionality of *K*_d_ is equal to that of 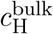. Solving Eq. (S33) yields the dependence of *ϕ*_b_ on buffer pH, as shown in Fig. S3. We can see that the relationship between *ϕ*_b_ and pH exhibits a monotonic behavior, indicating that the influence of the degree of ionization can be captured by pH.

**Figure S3:**
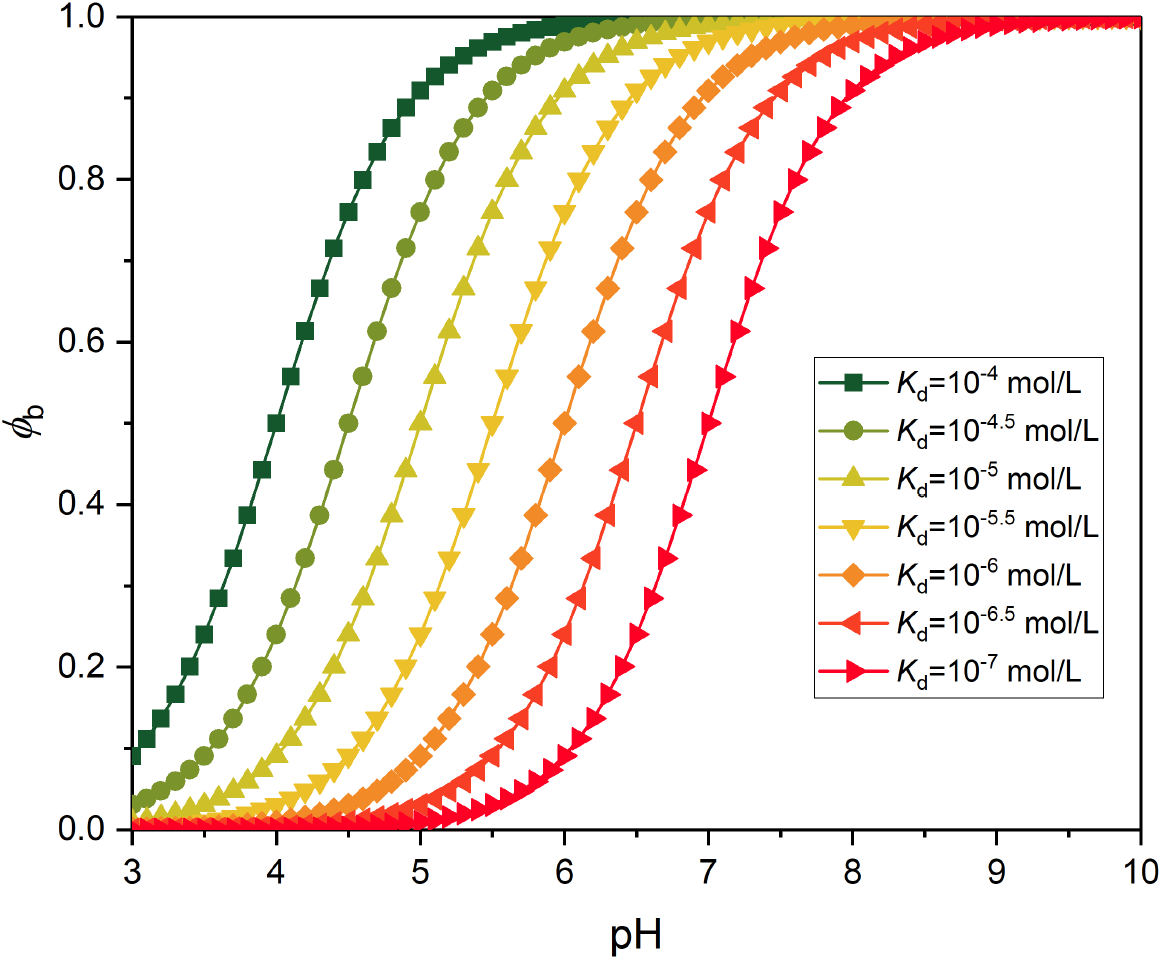
The dependence of *ϕ*_b_ on pH at different *K*_d_.

Moving forward, the cubic and higher terms in Eq. (S31) are neglected. Therefore, the sum of Eq. (S29), Eq. (S31), and Eq. (S32) yields the energy contribution of biopolymers

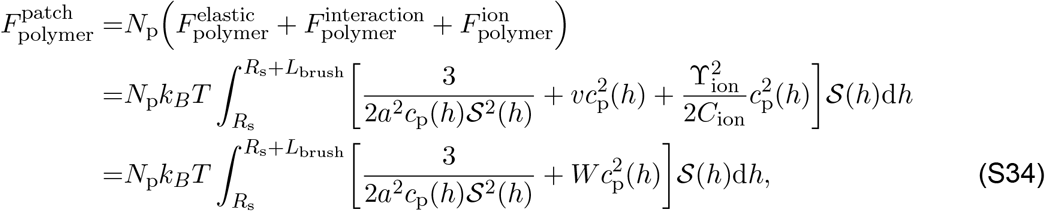

where *N*_p_ = *ρA*_0_ is the number of polymer chains grafted on the membrane, and 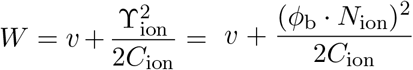. Here *A*_0_ stands for the membrane patch area covered by biopolymers and *ρ* is the grafting density. Plotting *W* as a function of *C*_ion_ and *ϕ*_b_ (Fig. S4) reveals that *W* decreases with increasing *C*_ion_ but increases with increasing *ϕ*_b_. Since *W* is given by the sum of the second virial coefficient *v* and 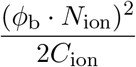, the trends observed in Fig. S4 imply that variations in ionic strength and the degree of ionization modify the effective monomer–monomer interaction coefficient, thereby regulating the effective excluded-volume interactions between monomers.

**Figure S4:**
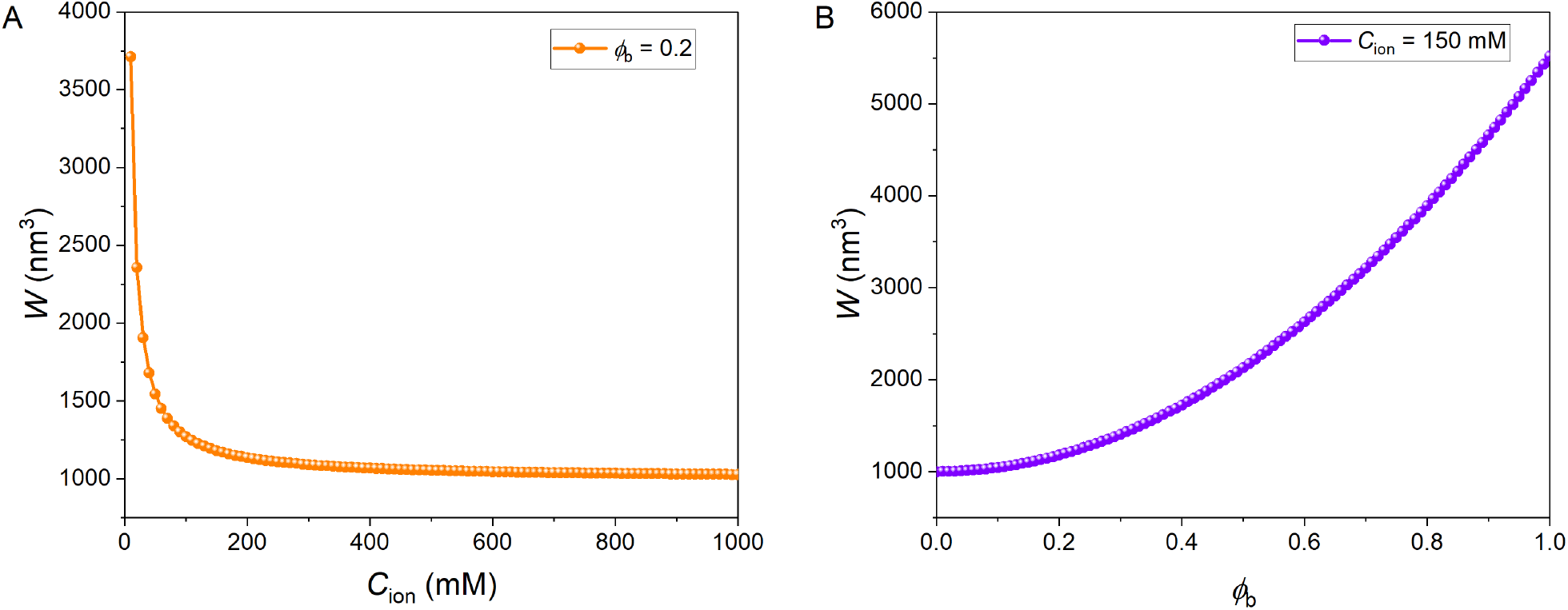
The dependence of *W* on *C*_ion_ and *ϕ*_b_.

In order to calculate the free energy, we need to further determine the local concentration of monomers *c*_p_(*h*) and the brush thickness *L*_brush_. On a planar membrane surface, the planar brush area per chain, *S* (*h*), is constant, i.e., *S* (*h*) = *S*. Minimizing the free energy of a planar brush *F*_polymer_ with respect to *c*_p_ by using 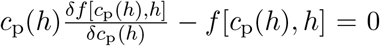 leads to the equilibrium polymer concentration 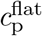 (see Ref. (*77*) for detailed steps)

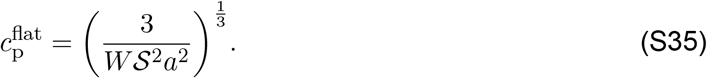

Here, 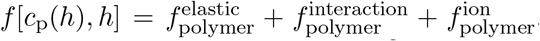. Using the conservation condition defined by Eq. (S28) yields the brush thickness 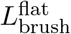 (see Ref. (*77*) for detailed steps)

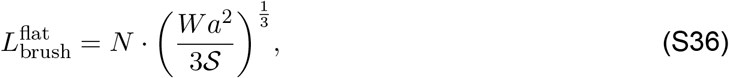

where we can see that the thickness of polymer brush is proportional to the total number of segments *N*. Hereafter, in our model, we use the number of segments, *N*, to capture the length of the polymer. When a polymer brush layer is anchored to a flat membrane surface, the effects of ionic strength and the degree of ionization on 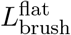 are shown in Fig. S5. Figures S5A and S5B demonstrate that the brush height decreases with increasing ionic strength but increases with increasing degree of ionization. This indicates that brush height is sensitive to changes in ionic strength and the degree of ionization on a flat membrane surface.

Substituting Eq. (S35) and Eq. (S36) into Eq. (S34) yields the energy contribution of biopolymers on a planar membrane surface

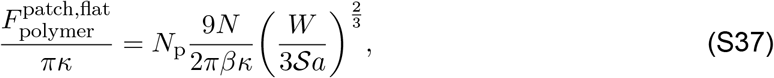

which corresponds to the case *α* = 0 in Eq. (3) in the main text. Thus, even for a flat membrane, the energy contribution by the biopolymers is directly proportional to the extent of grafting *N*_p_ and the length of the polymer brush *N*.

**Figure S5:**
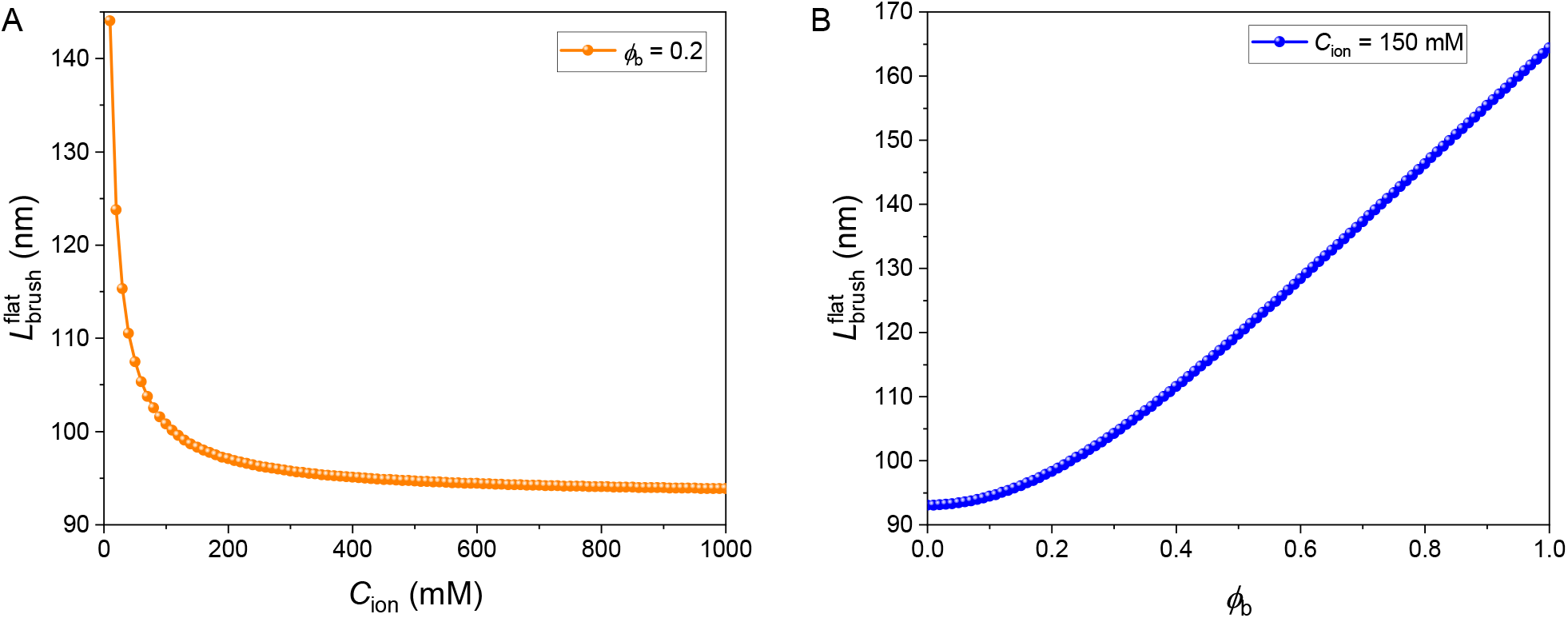
The dependence of 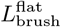 on *C*_ion_ and *ϕ*_b_.

On a spherical membrane surface, the corresponding monomer density profile *c*_p_(*h*) and brush thickness *L*_brush_ are, respectively, expressed as (see Ref. (*77*) for a detailed calculation)

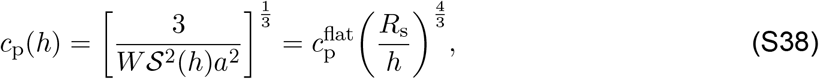

and

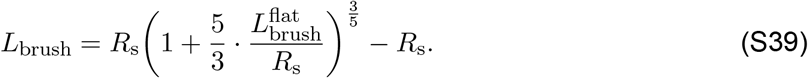

Thus, given the equilibrium membrane radius *R*_s_, Eq. (S39) can be applied to evaluate the height of the biopolymer brush layer. To further elucidate how ionic strength and the degree of ionization affect the brush height when the polymer brush layer is tethered to a curved membrane surface, we plot the dependence of *L*_brush_ on *C*_ion_ and *ϕ*_b_ by using Eq. (S39). Figures S6A and S6B show that decreasing *C*_ion_ and increasing *ϕ*_b_ lead to the swelling of the polymer brush on the curved membrane surfaces. This suggests that ionic strength and the degree of ionization play important roles in determining brush height even in the presence of membrane curvature.

Similarly, substituting Eq. (S38) and Eq. (S39) into Eq. (S34) leads to the energy contribution of biopolymers on a spherical membrane surface as

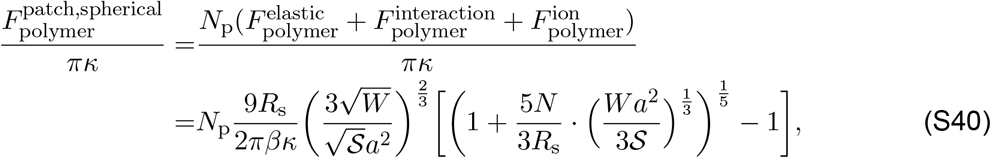

which corresponds to the case 0 *< α* ≤ *π* in Eq. (3) in the main text.

Combining Eqs. (S24), (S25), (S37), and (S40) yields Eq. (3) in the main text. In the following section, we derive the free energy associated with the free membrane region.

**Figure S6:**
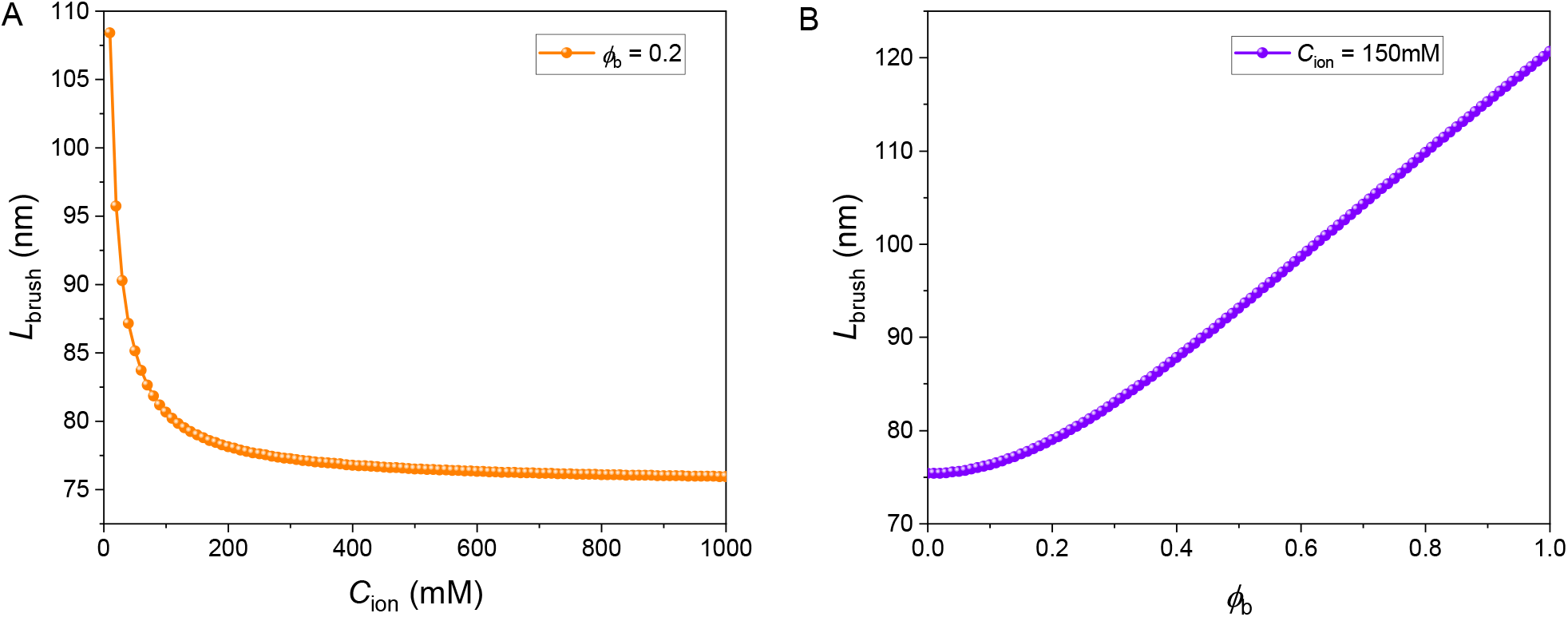
The dependence of *L*_brush_ on *C*_ion_ and *ϕ*_b_.

### 4 Energy contribution of the free part of the membrane

#### 4.1 Fundamental differential geometry of embedded surfaces

In order to derive the free energy functional of the free membrane part, we briefly outline the pre-requisite differential geometry quantities of a two-dimensional fluid membrane surface. Note that a detailed description of relations of differential geometry for the surface can be found in an extensive literature (e.g. Refs. (*81, 82*)). We modeled here the plasma membrane as a two-dimensional (2D) curved surface embedded in Euclidean three-dimensional (3D) space ℝ^3^. The surface of the membrane can be described by a parametrization ***X***(*u*^1^, *u*^2^), where *u*^1^ and *u*^2^ are the intrinsic curvilinear coordinates associated with a parametrization of the surface. At an arbitrary position of the membrane surface, the internal coordinate *u*^*i*^ defines a natural local covariant basis vector in the tangent plane of the surface given by

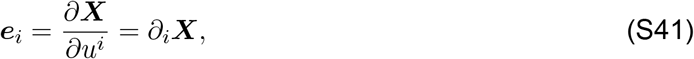

where the index *i* ∈ *{*1, 2*}*. In general, tangent vectors are neither orthogonal to each other nor of unit length. Using the tangent vector, the unit normal vector to the surface (i.e. ***n*** *·****e***_*i*_ = 0 and ***n*** *·****n*** = 1) is defined as

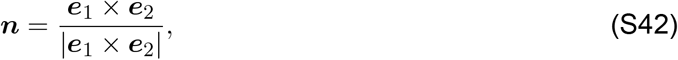

where × is the cross product, and |·| gives the normal of the vector. The set {***e***_1_, ***e***_2_, ***n***} comprising the tangent vectors and the extrinsic unit normal vector, forms a basis of the Euclidean 3D-space ℝ^3^. Lower indices correspond to covariant coordinates and upper indices to contravariant coordinates. Tangent vectors also define the metric tensor with its components are given by

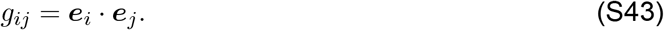

Note that the inverse of the metric tensor, *g*^*ij*^ = (*g*_*ij*_)^−1^, satisfies

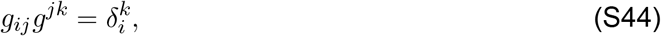

where Einstein summation convention is employed and 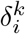 is the Kronecker delta. The metric tensor captures the distance information on the surface, being an intrinsic geometrical property of that surface. Note that the symmetric metric tensor (*g*_*ij*_ = *g*_*ji*_) can be used to raise and lower indices, respectively; for example,

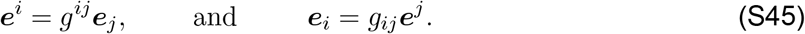

The metric tensor can be also exploited to calculate the infinitesimal area element 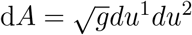, where *g* is the determinant of the metric tensor *g* = det(*g*_*ij*_). In addition, using the tangent vectors and their corresponding normal vector, the curvature tensor *K*_*ij*_ associated to ***X*** can be defined as

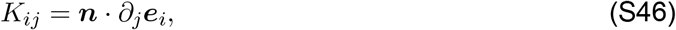

and 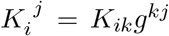. This tensor is an extrinsic curvature tensor which measures the bending of the 2D surface in the 3D space. 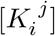 is also a symmetric matrix with two eigenvalues which are commonly called the principal curvatures. Hence the mean curvature *H* and the Gaussian curvature *K* are calculated as

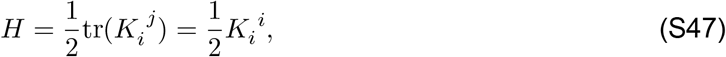

and

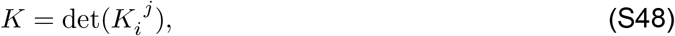

which describes the shape of the membrane surface due to its embedding in ℝ ^3^. The Gaussian curvature can also be written as *K* = det(*K*_*ij*_)*/*det(*g*_*ij*_).

In the following subsection, we employ the differential geometric formalism introduced above to parametrize the free membrane surface and derive the corresponding free energy.

#### 4.2 Arclength parametrization

**Figure S7:**
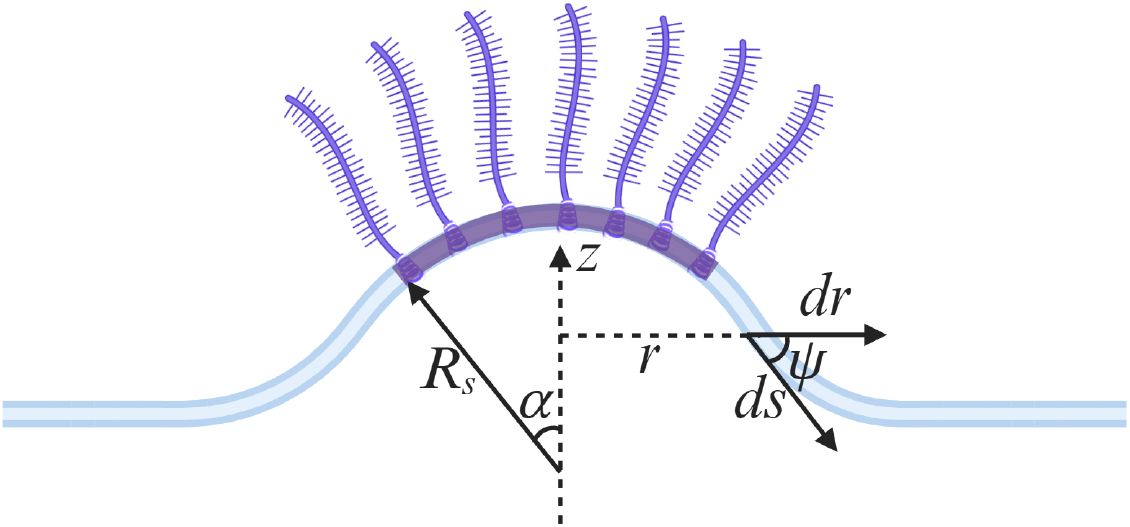
Schematic of a membrane consists of two parts: one patch domain composed of grafted polymers (purple), and the free membrane.

To compute the mechanical energy of the free membrane part, we assume that the membrane geometry is axisymmetric about the *z* axis. The surface function ***X*** of the free membrane part can be specified by a pair of internal curvilinear coordinates {*u*^1^, *u*^2^} which is associated with a parametrization of the free membrane surface. In cylindrical coordinates, defining *s* and *θ* as the arclength of the membrane and the azimuthal angle, the surface coordinates become ***X***(*u*^1^, *u*^2^) = ***X***(*s, θ*), and the surface is ***X***(*s, θ*) = [*r*(*s*) cos *θ, r*(*s*) sin *θ, z*(*s*)]. For the free part of the membrane, the arclength parametrization specifies the surface shape as a function of the arclength *s* along the contour of the surface profile and the azimuthal angle about the symmetry axis of the system under consideration. We denote the coordinate parallel to the axis of symmetry by *z*(*s*), the (inplane) coordinate perpendicular to the axis of symmetry by *r*(*s*), and the angle between the tangent vector ***e***_*s*_ = ∂_*s*_***X*** and the *r*-axis by *ψ*(*s*), as shown in Fig. S7. Then the elastic energy of the free membrane part comes from the axisymmetrically curved membrane shape can be described by *r*(*s*), *z*(*s*) and *ψ*(*s*), where *s* is the arc length of the free membrane. The coordinates *r*(*s*) and *z*(*s*) depend on *ψ*(*s*) through constraints 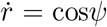 and 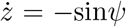, where the dots denote a derivative with respect to the arc length. Using Eq. (S43), one finds that the metric tensor is diagonal with *g*_*ss*_ = 1 and *g*_*θθ*_ = *r*^2^. Using Eq. (S47), the scalar invariant mean curvature *H* denotes the average of the two principal curvatures and is given by *H* = *g*^*ij*^*K*_*ij*_*/*2, where *g*^*ij*^ = (*g*_*ij*_)^−1^. The two principal curvatures are given by

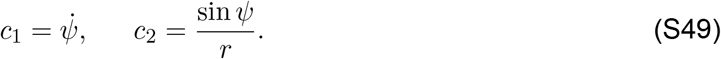

The elastic energy of the free membrane part then reads

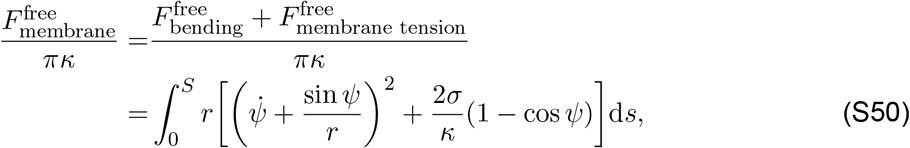

where dividing by *πκ* normalizes the energy. In addition, we introduce two Lagrangian multipliers *µ* and *ν*to impose the geometric constraints

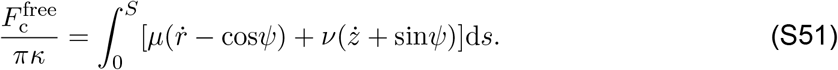

Therefore, the total free energy of the free membrane can be written as (*42, 43*)

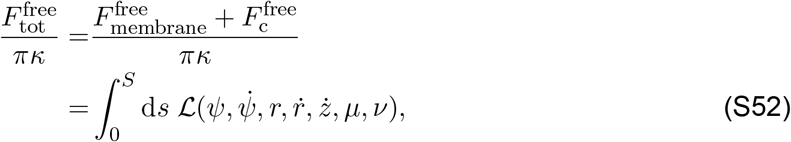

where ℒ is a Lagrangian defined by

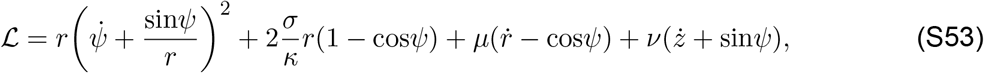

which corresponds to Eq. (4) in the main text. Here *µ*(*s*) and *ν*(*s*) are Lagrange multipliers to impose the geometric constraints between *r, z* and *ψ*. By introducing the variation: *ψ*(*s*) = *ψ*_0_(*s*) + *δψ*(*s*), *r*(*s*) = *r*_0_(*s*) + *δr*(*s*), *z*(*s*) = *z*_0_(*s*) + *δz*(*s*), *µ*(*s*) = *µ*_0_(*s*) + *δµ*(*s*), *ν*(*s*) = *ν*_0_(*s*) + *δν*(*s*), *S* = *S*_0_ + *δS*, then the variation of the functional 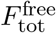 reads

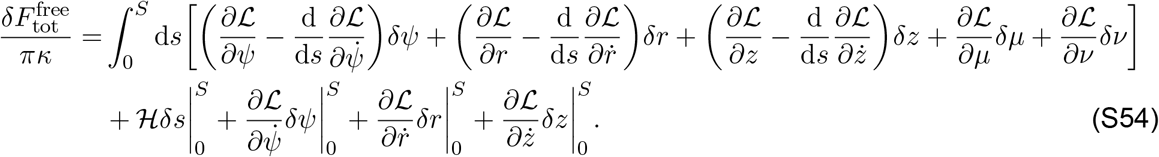

which contains both the bulk terms (the terms in the square bracket) and the boundary terms (last three terms). Here, ℋ is the Hamiltonian function. A set of Euler-Lagrange equations can be obtained by the vanishing bulk terms, which are reduced to

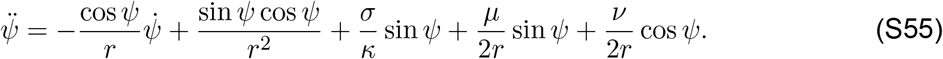

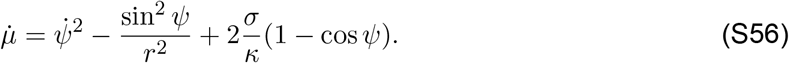

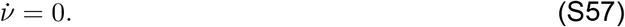

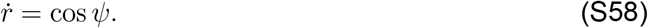

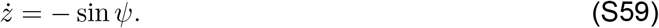

The stationary membrane shapes could be obtained by directly solving the corresponding Euler-Lagrange equations (S55)-(S59) subject to suitable boundary conditions. The arclength parametrization of surfaces provides an elegant approach for the representation of axisymmetric shapes with, potentially, large gradients. In the arclength parametrization, the Euler-Lagrange equations are, in general, highly nonlinear and must be solved numerically.

#### 4.3 Boundary conditions

For completeness, we need boundary conditions to numerically solve the above Euler-Lagrange equations. Here, we take the value of *ν*as a constant which equals zero, i.e., *ν*(0) = 0 due to the fact that its first order derivative is zero. With boundary conditions at the contact point between the boundary of the glycocalyx attached patch and the free membrane, *ψ*(0) = *α* and *r*(0) = *R*_*s*_ sin *α*, as well as at the infinity *ψ*(∞) = 0, and *z*(∞) = 0. In addition, we consider a homogeneous membrane, so that the Lagrangian *L* is explicitly independent of the arc length *s*. As a result, the Hamiltonian 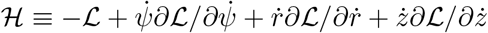 is a conserved quantity (*43*) given by

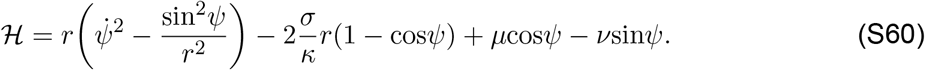

Due to that ℋ is conserved along the arc length, i.e., ℋ (*s*) = 0. We therefore impose the last boundary condition which is ℋ (0) = 0. We numerically solve the shape equations for various *α* and obtain the total free energy *F*_tot_(*α*) as a function of the opening angle *α*. The set of Euler-Lagrange equations plus its boundary conditions constitute a well-defined boundary value problem (BVP) that can be numerically solved by different methods. For example, ‘shooting’ method has been employed widely for numerically finding the stationary shapes implied by the arclength parametrization. However, here the ordinary differential equations along with the boundary conditions are solved in Matlab using ‘bvp5c’ solver.

By combining Eqs. (S55) and (S60), and letting *ν*= 0, one can derive the general shape equation

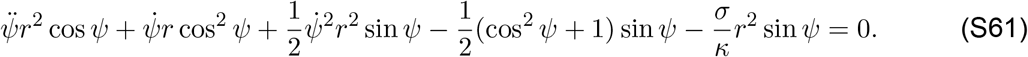

As long as the membrane equilibrium shape is determined, the total energy of the system can be obtained by summing Eq. (S24), Eq. (S25), Eq. (S37), Eq. (S40), and Eq. (S52). According to the model developed in this section, we numerically solve the shape equations for various *α* and obtain the total energy *F*_tot_(*α*) as a function of the opening angle *α*. Based on the optimal opening angle *α*_min_ obtained via minimizing the total energy, we can determine the degree of membrane bending.

## 5 Total free energy

In this work, we assume that the area of the polymer attached membrane patch is fixed as 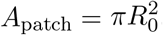, where *R*_0_ is the in-plane radius of the flat membrane. The requirement of the constant area of the polymer attached membrane patch gives rise to 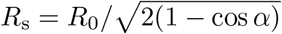 and the base radius 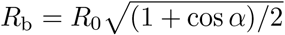. Hence, the total free energy is given by

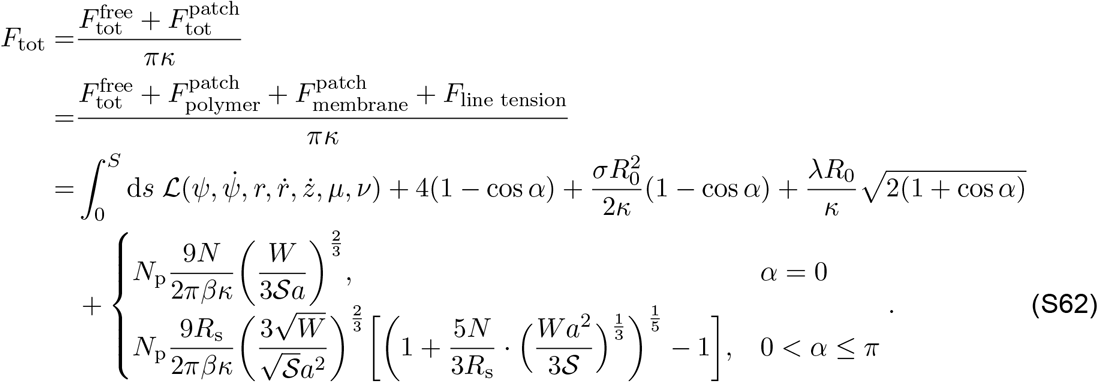

The corresponding boundary conditions for the free part of the membrane become *ψ*(0) = *α* and 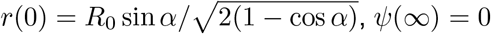, and *z*(∞) = 0.

Here, we demonstrate a simulation framework for calculating the total free energy of the system, with the flow chart shown in Fig. S8.

**Figure S8:**
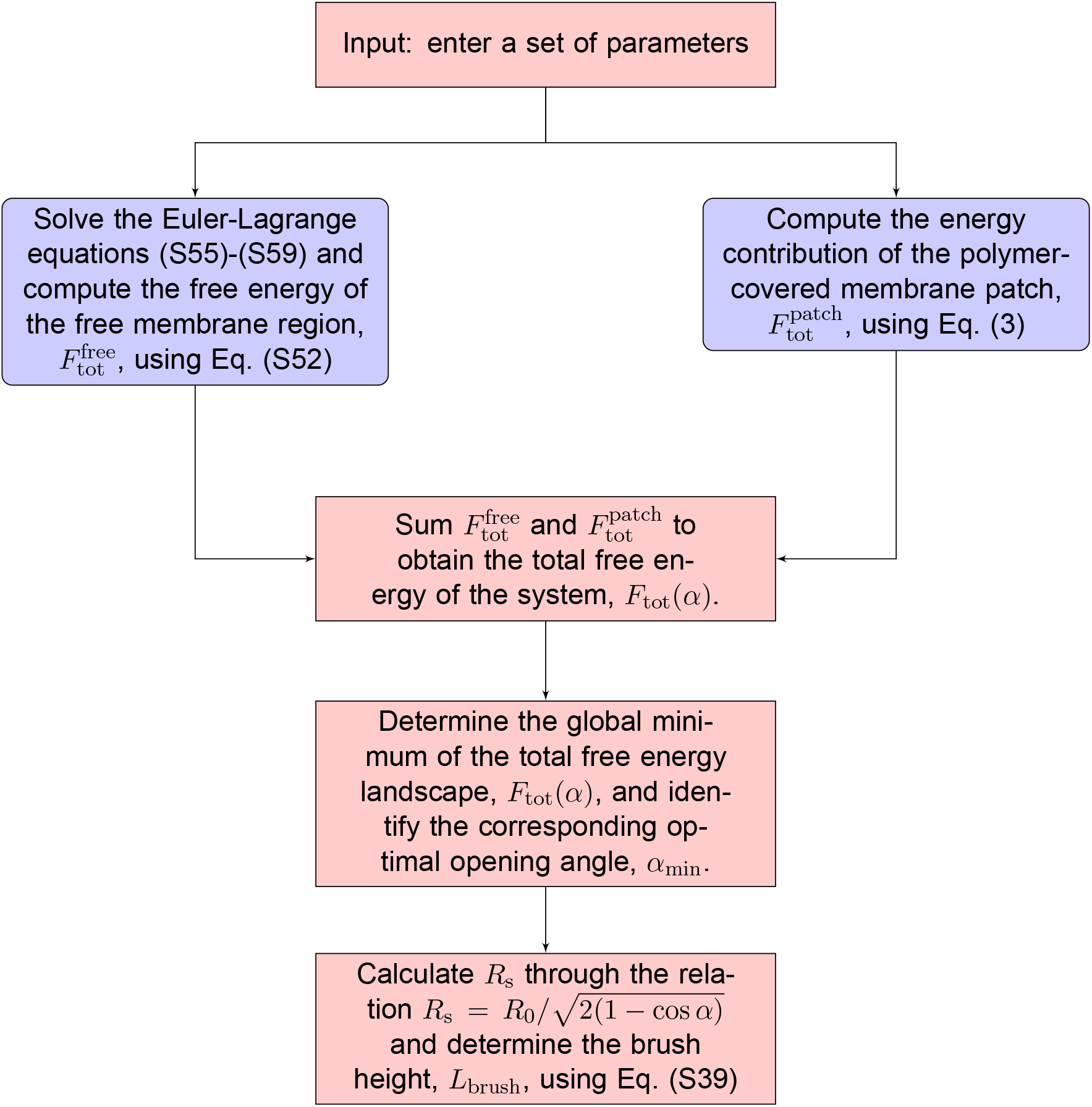
Simulation flow chart illustrating the procedure for determining the total free energy of the system at a fixed set of parameters. The default parameter values are listed in Table S1.

Consistent with the results shown in Fig. 6, the truncated his-AP180CTD-1/3 construct exhibited a similar sensitivity to ionic strength and pH, as shown in Fig. S9. Elevating the salt concentration from 50 mM to 500 mM suppressed outward tubule formation from 39.1% ± 5.6% to 25.1% ± 3.5% (*P* = 0.021), while increasing the buffer pH from 6.4 to 8.4 enhanced tubulation frequency from 24.4% ± 2.6% to 35.9% ± 2.4% (Fig. S9B,D; *P* = 0.047).

**Figure S9:**
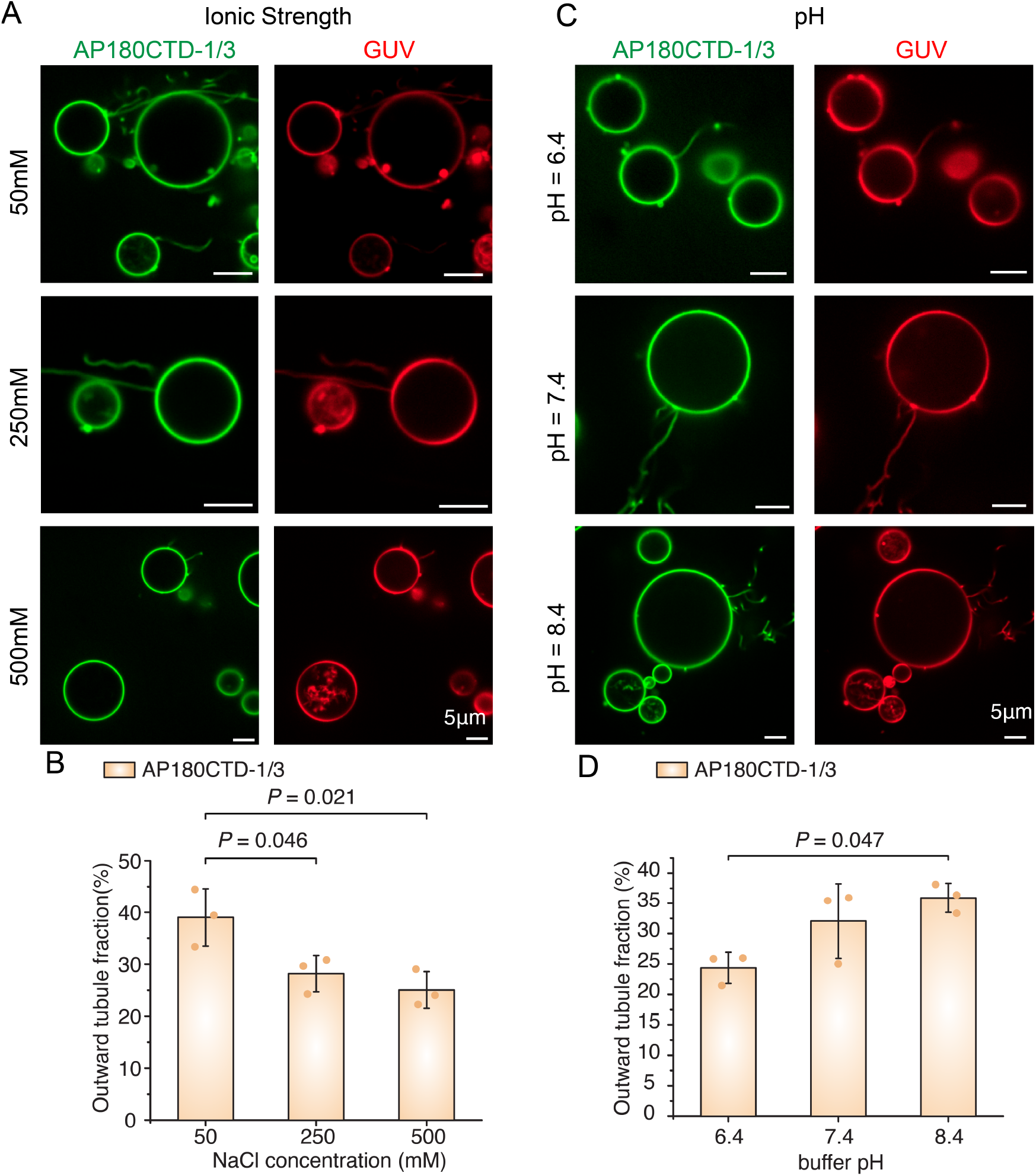
Effects of his-AP180CTD-1/3 on GUV tubulation. (A, C) Representative images and (B, D) quantification of outward tubule formation on GUVs incubated with 1 *µ*M 1/3 truncation of his-AP180CTD. Experiments were performed under varying NaCl concentrations (50–500 mM at pH 7.4) or varying pH levels (pH 6.4–8.4 at 150 mM NaCl). GUV lipid composition and image analysis were identical to those in Fig. 6. Error bars represent the SD of three independent trials.

### Critical conditions for membrane shape transition

Having established how intrinsic membrane-tethered polymer properties influence this critical threshold (orange solid line in Fig. 3E), we now turn to ask how solution conditions affect the critical grafting density, *ρ*_c_, that triggers the snap-through shape transition. To answer this question, we plot the dependence of the critical grafting density, *ρ*_c_, on *ϕ*_b_ and *C*_ion_ in Figs. S10A and S10B. Our results reveal that increasing the degree of ionization *ϕ*_b_ and reducing the ionic strength *C*_ion_ lowers the threshold *ρ*_c_ required to trigger the discontinuous transition in membrane structures. The inversely proportional relationship between *ρ*_c_ and *ϕ*_b_, and the proportional relationship between *ρ*_c_ and *C*_ion_ imply that the effects of *ϕ*_b_ and *C*_ion_ on the curvature generation are strikingly different. Overall, these observations suggest that longer polymers (with larger brush height) and more highly ionized polymers at lower ionic strength enhance curvature generation capability and thereby reduce the critical grafting density of the polymer required to trigger the formation of a spherical membrane structure.

**Figure S10:**
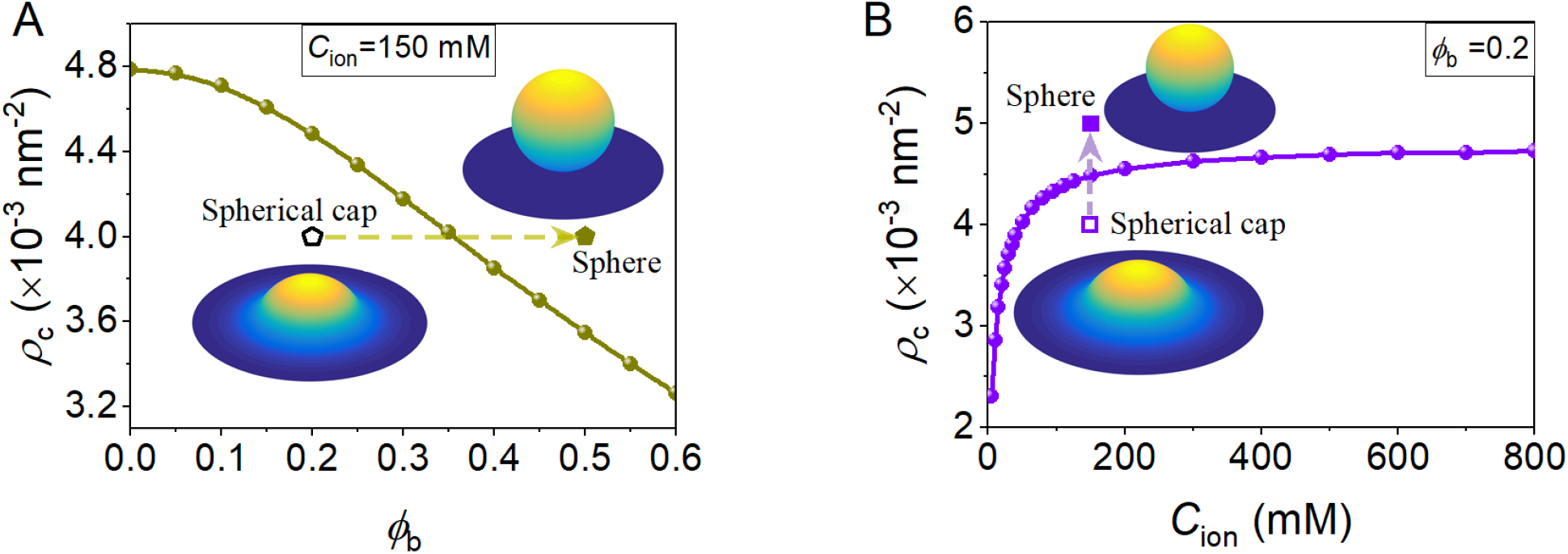
The dependence of critical grafting density, *ρ*_c_ on (A) *ϕ*_b_ and (B) *C*_ion_.

To illustrate the effect of grafting density on membrane shape, we plot the optimal opening angle, *α*_min_, and the spherical radius, *R*_s_, as a function of grafting density for different degrees of ionization in Figs. S11A and S11B.

**Figure S11:**
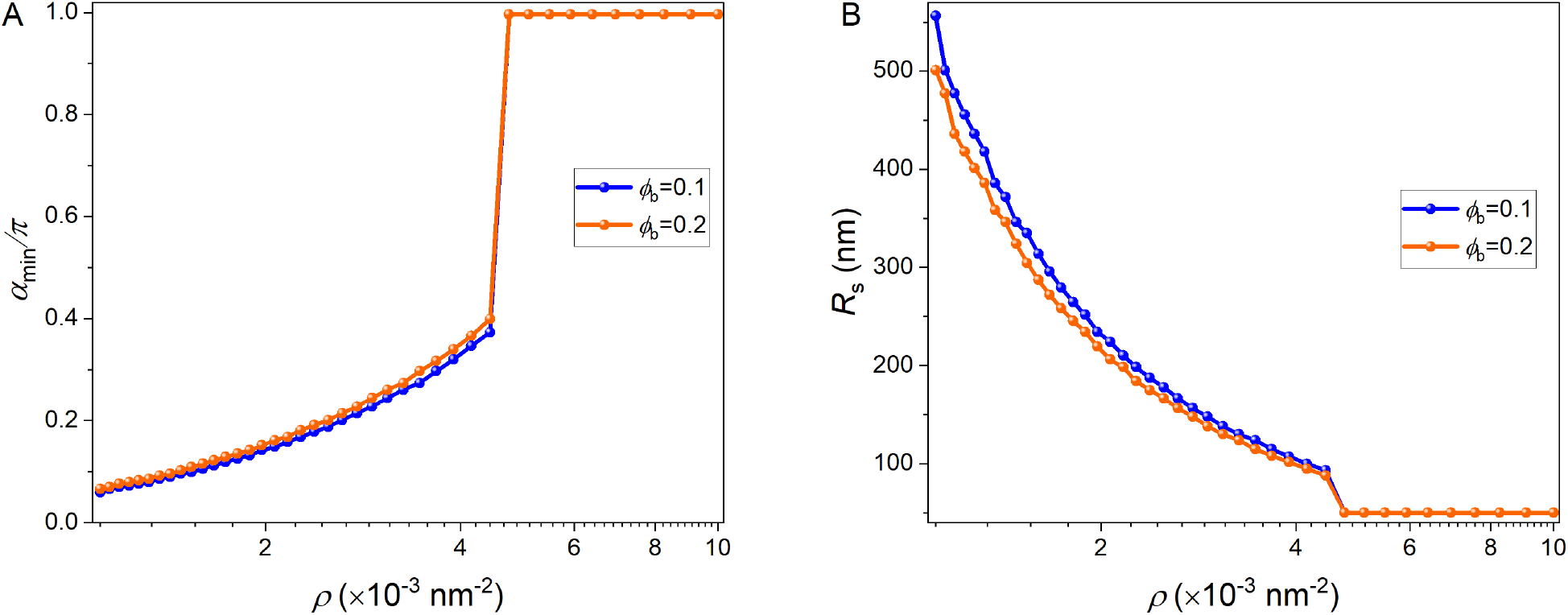
The dependence of (A) optimal opening angle *α*_min_ and (B) *R*_s_ on polymer grafting density.

#### 6 Numerical parameters

The numerical parameters used for the plots are given in Table S1.

**Table S1:** Parameters used in our numerical estimates.

| Parameter | Meaning | Estimated range | References | Default value |
| --- | --- | --- | --- | --- |
| $\kappa$ | Bending rigidity | 10–400 $k_B T$ | (83–86) | 20 $k_B T$ |
| $\sigma$ | Membrane tension | $10^{-7}$–$10^{-3}$ N/m | (40, 84, 87, 88) | 0.012 $k_B T/\text{nm}^2$ |
| $\lambda$ | Line tension | 0–100 pN | (84, 89, 90) | 0 $k_B T/\text{nm}$ |
| $R_0$ | Patch radius | 20–100 nm | (32, 84, 91–93) | 100 nm |
| $\xi$ | Grafting distance | 10–100 nm | (4, 20) | 18.2 nm |
| $a$ | Effective monomeric segment size in the biopolymer | 10–20 nm | (2, 4, 32, 88, 94, 95) | 10 nm |
| $a_m$ | Monomer size | $\sim 0.35$ nm | (2, 4, 32, 94, 95) | 0.35 nm |
| $N$ | Total number of monomeric segments in a brush | 10–50 | (2, 4, 32, 95) | 20 |
| $\phi_b$ | Degree of ionization | 0–1 | (4, 44) | 0.2 |
| $C_{\text{ion}}$ | Ionic concentration | 5–1000 mM | (4, 44) | 150 mM |
| $A_p$ | Polymer area | $\sim 100$ nm<sup>2</sup> | (96) | 100 nm<sup>2</sup> |
| $\omega$ | Polymer-membrane interaction | $\sim -10$–$10$ $k_B T$ | (16, 97) | 3.75 $k_B T$ |
| $\bar{\chi}$ | Polymer-polymer interaction | $\sim -10$–$10$ $k_B T$ | (98) | $-7.5$ $k_B T$ |
| $e$ | Elementary charge | $1.6 \times 10^{-19}$ C | – | $1.6 \times 10^{-19}$ C |
| $\epsilon_0$ | Vacuum permittivity | $8.854 \times 10^{-12}$ F m<sup>-1</sup> | – | $8.854 \times 10^{-12}$ F m<sup>-1</sup> |
| $\epsilon_r$ | Relative permittivity of water | 80 | – | 80 |
| $k_B T = 1/\beta$ | Thermal energy | $4.1 \times 10^{-21}$ J | – | $4.1 \times 10^{-21}$ J |

**Table S2:** Notation and list of symbols.

| Notation | Description |
| --- | --- |
| $\kappa$ | Bending rigidity |
| $\sigma$ | Membrane tension |
| $H$ | Mean curvature |
| $c_1, c_2$ | Principal curvatures of the membrane |
| $c_{\text{eff}}$ | Effective spontaneous curvature of the membrane |
| $\phi_V, \phi_t$ | Area fraction of polymers on the planar cell surface and on the membrane tube |
| $A_p$ | Polymer area |
| $\omega$ | Polymer-membrane interaction |
| $\chi$ | Polymer-polymer interaction coefficient |
| $\lambda_D$ | Debye screening length |
| $l_B$ | Bjerrum length |
| $l_m$ | Membrane thickness |
| $L_t$ | Length of membrane tube |
| $R_t, R_V$ | Radius of membrane tube and vesicle |
| $\Delta p$ | Hydrostatic pressure difference across the cell membrane |
| $e$ | Element charge |
| $\epsilon_0$ | Vacuum permittivity |
| $\epsilon_r$ | Relative permittivity of water |
| $\alpha$ | Opening angle |
| $R_0$ | Patch radius |
| $R_s$ | Radius of the spherical cap |
| $L_{\text{brush}}$ | The height of the brush |
| $\mathcal{S}$ | The local area per chain |
| $c_p$ | Local density profile of monomers |
| $v$ | The second virial coefficient |
| $w$ | The third virial coefficient |
| $\lambda$ | Line tension |
| $\xi$ | Grafting distance |
| $a$ | Effective monomeric segment size in the biopolymer |
| $a_m$ | Monomer size |
| $N$ | Number of effective monomeric segments in a brush |
| $N_{\text{ion}}$ | Number of ionizable sites per segment |
| $\Upsilon_{\text{ion}}, \phi_b$ | The degree of ionization of each segment, and of each amino acids |
| $c_s, C_{\text{ion}}$ | Salt concentration (Ionic strength) |
| $W$ | Effective monomer-monomer interaction coefficient |

| *continued from previous page* |  |
| --- | --- |
| $N_p$ | Number of polymer chains |
| $\rho$ | Grafting density |
| $A_0$ | Membrane patch area covered by biopolymers |
| $s$ | Arclength |
| $r$ | The coordinate perpendicular to the axis of symmetry |
| $z$ | The coordinate parallel to the axis of symmetry |
| $\psi$ | Angle made by surface tangent with the horizontal direction |
| $\mu, \nu$ | Lagrange multipliers |
| $\mathcal{L}$ | Lagrangian |
| $\mathcal{H}$ | Hamiltonian function |

#### 7 Experimental materials and methods

#### 7.1 Reagents

1-palmitoyl-2-oleoyl-sn-glycero-3-phosphocholine (POPC) and 1,2-dioleoyl-sn-glycero-3-{[N-(5-amino-1-carboxypentyl)iminodiacetic acid]succinyl} nickel salt (DGS-NTA-Ni) were obtained from Avanti Polar Lipids. Texas Red–DHPE, NeutrAvidin, tris-HCl, HEPES, isopropyl-*β*-D-thiogalactopyranoside (IPTG), *β*-mercaptoethanol, tris(2-carboxyethyl)phosphine (TCEP), and Triton X-100 were purchased from Thermo Fisher Scientific. Tryptone, yeast extract, sodium chloride, sodium phosphate salts, urea, sodium tetraborate, EDTA, calcium chloride, glycerol, imidazole, phenylmethanesulfonyl fluoride (PMSF), poly-L-lysine (PLL), and Atto 488 NHS ester were purchased from Sigma-Aldrich. EDTA-free protease inhibitor tablets were obtained from Roche. DP-EG10-biotin was generously provided by D. Sasaki (Sandia National Laboratories, Livermore, CA) (*99*). Methoxy polyethylene glycol succinimidyl valerate (mPEG-SVA, MW 5000) and biotin-PEG-SVA (MW 5000) were purchased from Laysan Bio. All reagents were used as received without further purification.

### 7.2 Plasmid Construction

The plasmid encoding AP180CTD (rat AP180 residues 328–898) in the pET32c vector was kindly provided by E. Ungewickell (Hannover Medical School, Germany). Histidine-tagged AP180CTD (his-AP180CTD) was subcloned into a pGEX-4T-2 vector to introduce an N-terminal glutathione S-transferase (GST) tag for enhanced protein stability during purification, as previously described (*100*). A truncated AP180CTD construct (his-AP180CTD-1/3) was generated by insertion of a stop codon at residue A213. All constructs were verified by DNA sequencing.

### 7.3 Protein expression and purification

His-AP180CTD and his-AP180CTD-1/3 were expressed in Escherichia coli BL21Star(DE3) pLysS cells (NEB, catalog no. C2530). Transformed cells were cultured at 30^*°*^C to an OD600 of approximately 0.8, followed by induction with 1 mM IPTG for 24 h at 12^*°*^C with shaking at 200 rpm. All purification steps were performed at 4^*°*^C. Cells from 2 L cultures were harvested by centrifugation at 4,785 × g for 20 min and resuspended in lysis buffer containing 0.5 M tris-HCl (pH 8.0), 5% glycerol, 5 mM EDTA, 5 mM TCEP, 1 mM PMSF, EDTA-free protease inhibitor tablets, and 1% Triton X-100. Cell suspensions were homogenized using a Dounce homogenizer and lysed by sonication on ice. Lysates were clarified by centrifugation at 26,581 × g for 25 min. The supernatant was applied to a Glutathione Sepharose 4B affinity column (Cytiva, catalog no. 17075605) and washed extensively with lysis buffer containing 0.2% Triton X-100, followed by detergent-free lysis buffer. Bound proteins were eluted using lysis buffer supplemented with 15 mM reduced glutathione and subsequently buffer-exchanged into 50 mM tris-HCl (pH 8.0), 10 mM CaCl2, 150 mM NaCl, and 5 mM EDTA using Zeba desalting columns (Thermo Fisher Scientific, catalog no. 89891). GST tags were removed by thrombin cleavage using the Thrombin CleanCleave Kit (Sigma-Aldrich, catalog no. RECOMT) for 14 h at 4^*°*^C with gentle agitation. Cleaved GST and uncleaved proteins were removed by a second Glutathione Sepharose 4B purification step. Purified proteins were concentrated using Amicon Ultra centrifugal filters (MilliporeSigma, catalog no. UFC903024), flash-frozen in liquid nitrogen, and stored at −80^*°*^C.

### 7.4 Protein labeling

Proteins were fluorescently labeled with Atto 488 NHS ester. Labeling reactions were performed in 25 mM HEPES, 150 mM NaCl, and 5 mM TCEP (pH 7.4). Dye was added at a twofold molar excess relative to protein and incubated for 30 min at room temperature, yielding labeling efficiencies of approximately 0.8–1.5 fluorophores per protein molecule. Labeled proteins were separated from free dye and buffer-exchanged into storage buffer containing 50 mM tris-HCl (pH 8.0), 10 mM CaCl2, 150 mM NaCl, 5 mM EDTA, and 5 mM TCEP using 3 kDa molecular weight cutoff Amicon filters. Protein and dye concentrations were quantified by UV-visible spectroscopy. Aliquots were flash-frozen in liquid nitrogen and stored at −80^*°*^C. Unless otherwise noted, experiments using fluorescently labeled protein were performed with a mixture containing 10% labeled and 90% unlabeled protein.

### 7.5 GUV preparation

GUVs were composed of 83 mol% POPC, 15 mol% DGS-NTA-Ni, and 2 mol% DP-EG10-biotin. For fluorescence imaging, 0.1 mol% Texas Red–DHPE was included. GUVs were prepared by electroformation as previously described (*101*). Lipid mixtures dissolved in chloroform were deposited onto indium tin oxide (ITO)-coated glass slides and dried under vacuum for at least 2 h. Electroformation was performed at 55^*°*^C in glucose solutions with osmolarities matched to the corresponding experimental buffer conditions. Specifically, 170, 360, and 940 mOsm glucose solutions were used for buffers containing 50, 150, and 500 mM NaCl, respectively. Voltage was gradually increased from 50 to 1400 mV peak-to-peak over 30 min at 10 Hz, maintained at 1400 mV for 120 min, and finally increased to 2200 mV for an additional 30 min at 5 Hz. Prepared GUVs were stored at 4^*°*^C and used within 3 days.

### 7.6 GUV tethering and sample preparation

GUVs were tethered to glass coverslips for imaging as previously described (*17*). Briefly, glass coverslips were passivated with biotinylated PLL-PEG to minimize nonspecific adsorption and facilitate GUV tethering. PLL-PEG was synthesized by reacting PLL with a mixture of mPEG-SVA and biotin-PEG-SVA at a molar ratio of 98:2. The reaction was carried out in 50 mM sodium tetraborate buffer (pH 8.5) with a lysine-to-PEG molar ratio of 5:1 for 6 h at room temperature. Excess reactants were removed by desalting into 25 mM HEPES and 150 mM NaCl (pH 7.4). Imaging chambers were assembled using silicone gaskets mounted on cleaned No. 1.5 glass coverslips. Coverslips and gaskets were cleaned with 2% Hellmanex III solution, rinsed thoroughly with water, and dried under nitrogen. PLL-PEG solution was added to each imaging well and incubated for 20 min, followed by extensive rinsing with experimental buffer. NeutrAvidin (4 μg per well) was then added and incubated for 10 min before rinsing. GUVs were diluted 1:13 in osmolarity-matched buffer and incubated in the wells for 10 min to allow tethering via biotin–NeutrAvidin interactions. Excess vesicles were removed by gentle rinsing prior to imaging. All experimental buffers contained 25 mM HEPES (pH 7.4) with the indicated NaCl concentration. Osmolarity was adjusted using glucose and verified using a vapor pressure osmometer.

### 7.7 GUV fluorescence imaging

Fluorescence imaging was performed using an Olympus SpinSR10 spinning-disk confocal super-resolution microscope equipped with a 100× oil immersion objective (NA 1.49). Excitation wave-lengths of 488 and 561 nm were used for imaging fluorescently labeled proteins and membrane dyes, respectively. Z-stack images were acquired at 0.5 μm intervals perpendicular to the membrane plane immediately after GUV tethering and following protein addition. Imaging was typically initiated 5 min after protein addition to allow membrane binding and equilibration. At least 30 randomly selected fields of view were analyzed for each experimental condition.

### 7.8 Statistical analysis

All GUV experiments were independently repeated at least three times. Image analysis was performed using ImageJ software, with more than 100 GUVs analyzed for each condition. Statistical comparisons between groups were performed using unpaired Student’s t-tests. Error bars represent standard deviation (SD), as indicated in the corresponding figure legends.

